# Mechanisms of traditional Chinese medicine in modulating cardiac microvascular endothelial cells in various injury models: A comprehensive systematic review

**DOI:** 10.1101/2024.09.05.611361

**Authors:** Huiwen Zhou, Hongxu Liu, Xiang Li, Juju Shang, Jiaping Chen, Huiqi Zong

**Author notes:** Corresponding author: (HL); (XL).

## Abstract

**Background:** The structural and functional failure of cardiac microvascular endothelial cells (CMECs) is a primary contributor to coronary microvascular dysfunction (CMD). Traditional Chinese medicine (TCM) has been identified as a potential therapeutic approach for preserving CMECs and mitigating CMD.

**Objective:** This systematic review aims to present the latest evidence on TCM intervention mechanisms in CMECs under diverse injury models.

**Methods:** This systematic review was performed following the parameters of the PRISMA statement (Preferred Reporting Items for Systematic Reviews and Meta-Analysis). A comprehensive literature search was conducted using PubMed, Embase, Web of Science, Scopus, China National Knowledge Infrastructure and China Biology Medicine disc. Reference lists of selected articles were reviewed to identify relevant studies. The search was not limited by year and was conducted solely in English. Eligible studies comprised publications describing in vitro studies that presented the latest evidence on TCM intervention mechanisms in CMECs under diverse injury models.

**Results:** A total of 63 papers were included in this study. According to the cell processing approach, 19 studies on ischemia or hypoxic injury models, 16 studies on Ischemia/reperfusion (I/R) or hypoxia/reoxygenation (H/R) injury models, 10 studies on inflammatory injury models, 5 studies on metabolic injury models, 3 studies on angiotensin II injury models, and 10 studies on other models. TCM exhibits structural and functional intervention capabilities in diverse damage conditions of CMECs. Its mechanism of action involves antioxidant, anti-apoptotic, anti-inflammatory effects, as well as regulation of energy metabolism through signaling pathways such as HIF-1α/VEGF, PI3K/AKT, MAPK, and NF-κB.

**Conclusions:** The CCM and its constituents modulate CMECs through multiple signaling pathways in response to various injury models, thereby conferring protection on the coronary microcirculation.

**Graphical Abstract:** 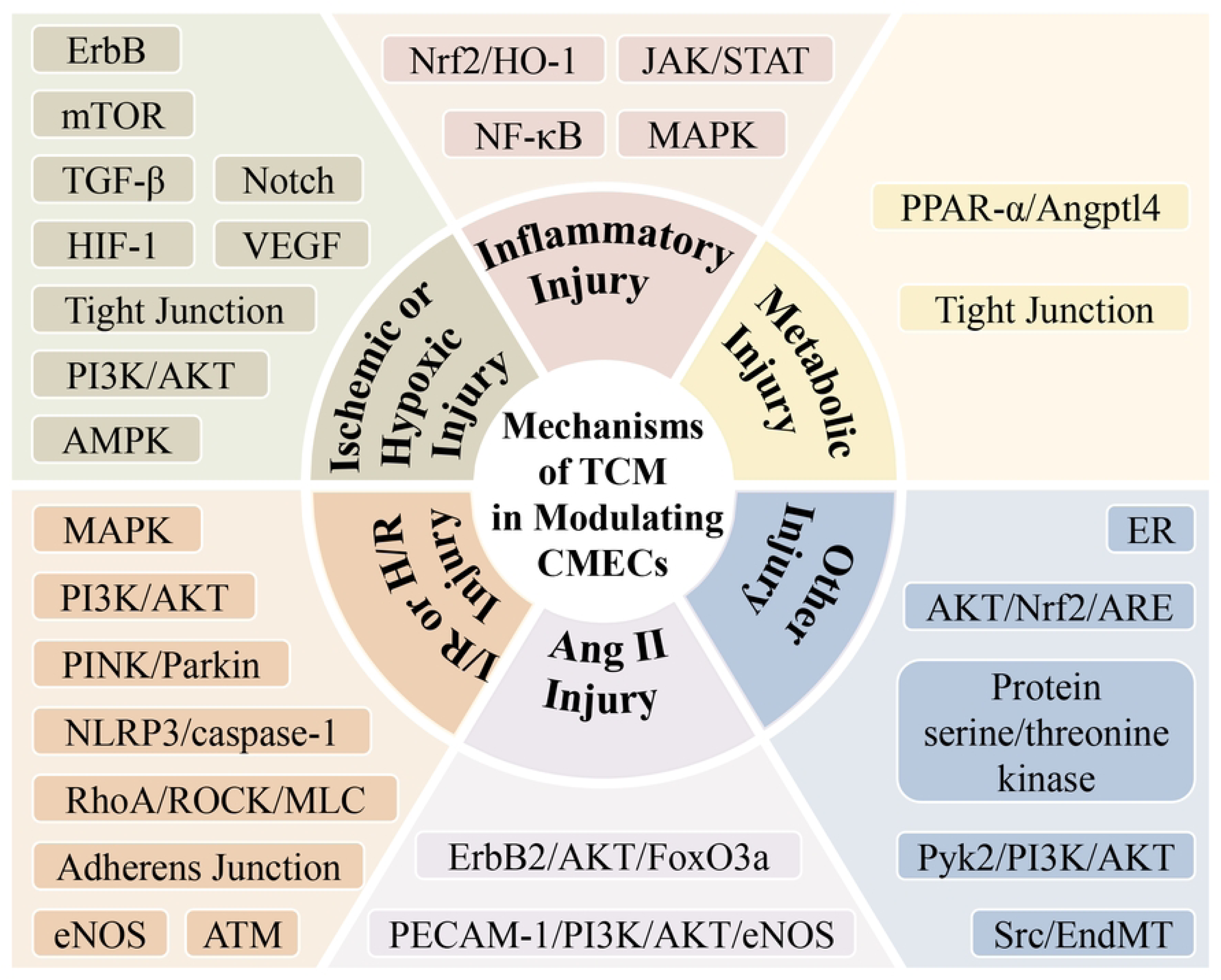

## 1. Introduction

Coronary microvascular disease (CMVD) is a clinical condition characterized by objective evidence of exertional angina and/or myocardial ischemia resulting from structural and/or functional abnormalities in the coronary microcirculation, triggered by various etiological factors. Coronary microvascular dysfunction (CMD) represents a crucial mechanism underlying CMVD [1]. CMD is implicated in different stages of cardiovascular disease and serves as a significant adverse prognostic factor for patients with ischemic heart disease (IHD) [2]. In recent years, CMD has garnered increasing attention due to its clinical relevance. Notwithstanding the high prevalence of CMD, effective treatment remains elusive. Therapeutic interventions such as nicorandil, statins, and angiotensin-converting enzyme inhibitors may offer potential benefits [3, 4]. However, the complete clinical picture remains incompletely understood. Traditional Chinese medicine (TCM), which has been utilized in clinical practice throughout China for nearly two millennia, has shown promise as a clinically viable approach for treating CMD [5, 6].

The development of CMD is significantly influenced by structural and/or functional abnormalities in cardiac microvascular endothelial cells (CMECs) [7]. In a physiological setting, the coronary microcirculation serves as the primary resistance artery in the coronary arteries. And it plays a crucial role in regulating coronary blood flow (CBF) [8]. CMECs, which constitute approximately one-third of all heart cells [9], are essential components of the coronary microcirculation [7], They play a critical role in controlling CBF and maintaining proper coronary microvascular function [10, 11]. When stimulated by pathological factors, CMECs lose their ability to proliferate, adhere, migrate normally or undergo apoptosis and secrete substances as usual. This can lead to abnormal contraction and diastolic function in microvessels as well as compromised integrity of the microvascular barrier and thinning of coronary microarterioles [12, 13]. Ultimately, this may result in reduced coronary flow reserve (CFR) and inadequate myocardial blood supply [14]. Therefore, investigating the mechanism through which compound Chinese medicine (CCM) and its constituents intervene in CMECs under various injury models can provide valuable insights into the potential of TCM for treating CMD. This review aims to present the latest evidence on TCM intervention mechanisms in CMECs under diverse injury models.

## 2. Methods

The systematic literature review was conducted in accordance with the guidelines outlined by the Preferred Reporting Items for Systematic Reviews and Meta-Analyses (PRISMA) statement [15] (S1 Table). The systematic review has been registered in the INPLASY platform for prospective registration with the registration number INPLASY202470092. Protocol details are available at <u>INPLASY Protocol 6561 – INPLASY</u>. Ethical approval was not required due to no human subjects being involved in this study.

### 2.1. Search Strategy

From database construction until March 2024, the electronic databases utilized for literature search encompassed PubMed, Embase, Web of Science, Scopus, China National Knowledge Infrastructure and China Biology Medicine disc. The keywords employed were “Chinese herbal medicine monomer”, “Chinese herbal medicine components”, “Chinese herbal compound”, “traditional Chinese medicine”, “herbal medicine”, “Chinese herbal medicine”, “Chinese herb”, “CHM”, “TCM”, “China Chinese herbal medicine”, “China extract”, “China fraction”, “China formula”, “China prescription, “CMECs”, “CMEC”, “coronary microvascular endothelial cells”, “cardiac microvascular endothelial cells”, “myocardiac microvascular endothelial cells”. The search scope was limited to full-text articles without any additional restrictions.

### 2.2. Flowchart Sketch of the Screening Process

The flowchart illustrating the systematic review screening process is presented in Fig 1. Initially, a total of 296 literature sources were identified and evaluated. Following a de-weighting procedure, 84 papers were excluded, while 212 papers met the inclusion criteria. The inclusion criteria consisted of the following: 1) the literature had to be written in English; 2) it had to be the full text; 3) the content of the literature had to be related to Chinese herbal medicines (CHM); 4) the study design had to involve CMECs or cardiac microvasculature. On the other hand, the exclusion criteria encompassed irrelevant literature, reviews, meta-analyses, case reports, conference proceedings, book chapters, letters to the editor, oral presentations, posters, and editorials. During the initial screening, 128 papers were excluded based on a skim of the title or abstract, leaving 84 papers for further evaluation. Among these, 12 papers lacked full text in English, 7 papers did not involve CMECs or cardiac microvessels in their design, and 2 papers were not focused on heart-related research. These 21 papers were consequently excluded. Finally, a total of 63 relevant studies were included in this review. Among them, 19 studies employed models of ischemic or hypoxic injury, 16 studies utilized models of ischemia/reperfusion (I/R) or hypoxia/reoxygenation (H/R) injury, 10 studies employed models of inflammatory injury, and 18 studies employed models of other types of injury, based on cellular processing methods (S2 Table). This review aims to comprehensively report and critically analyze the relevant studies. However, it should be noted that a meta-analysis was not conducted.

**Fig 1.**
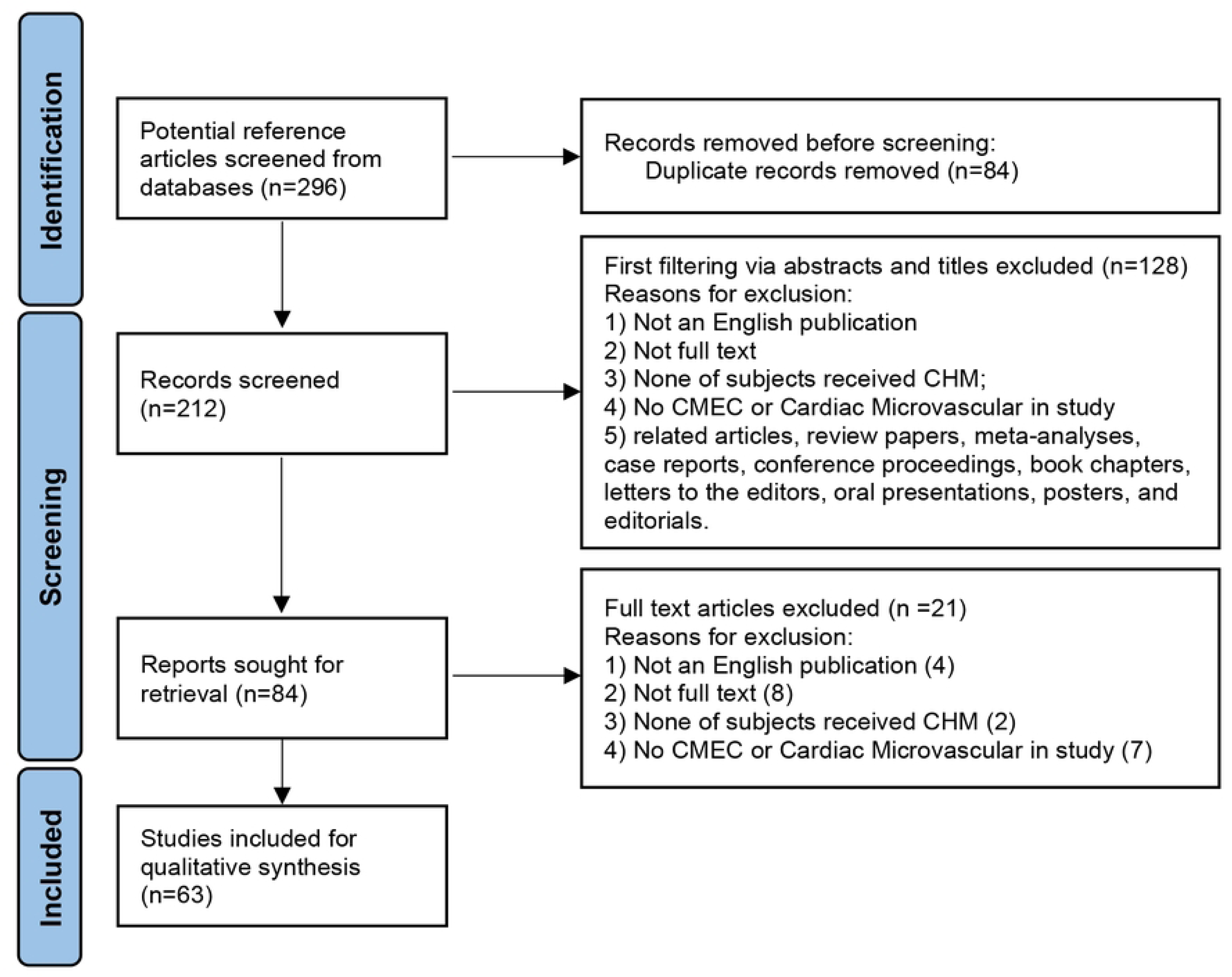
PRISMA flow chart.

## 3. Results

Among the 64 papers included, the intervention methods employed for CMECs exhibited considerable variation. Based on the specific cellular treatments applied, the literature can be categorized into distinct injury models, including ischemic or hypoxic injury, I/R or H/R injury, inflammatory injury, metabolic injury, angiotensin II (Ang II) injury, as well as other treatments. In these studies, a total of 16 CCMs and 18 major bioactive components (MBCs) were included. The CCMs and MBCs that are effective in regulating CMECs under various injury models are listed in S3 and S4 Tables. In the subsequent paragraphs, we will delve into a comprehensive review of the roles played by CCM and its constituents in different injury models, along with the underlying pathway mechanisms implicated.

### 3.1. TCM that Modify CMECs in Ischemic or Hypoxic Injury Model

Ischemia- and hypoxia-induced injury is widely regarded as the underlying pathological basis and initial process of numerous cardiovascular conditions. Protecting CMECs from injury caused by ischemia and hypoxia represents a crucial therapeutic approach for addressing a range of cardiovascular diseases [16]. Studies have discovered that TCM compounds or core constituents in regulating this injury plays a certain influence. and the involved mechanisms are illustrated in Fig 2 and summarized in Table 1.

**Fig 2.**
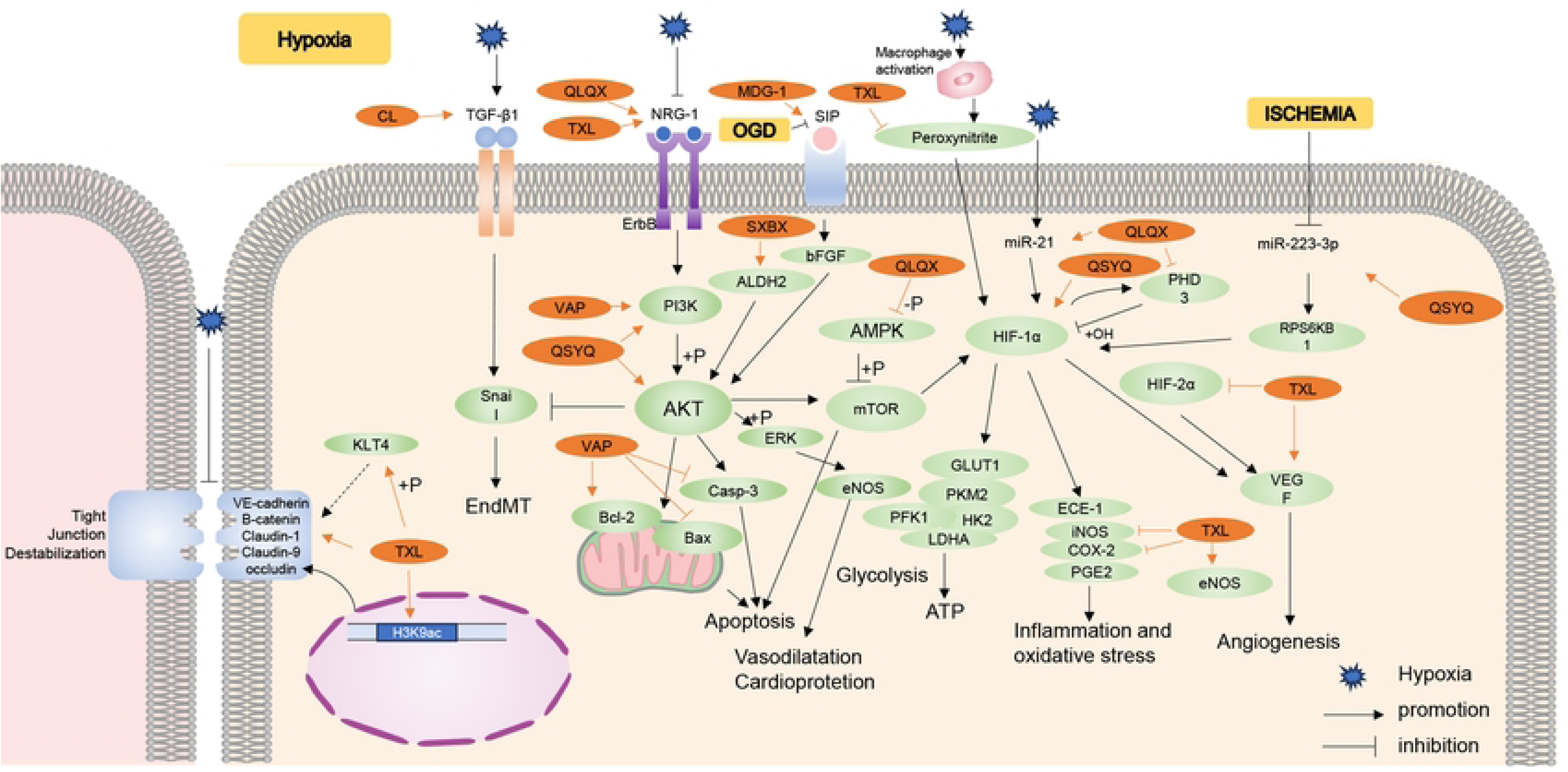
TCM regulate various signaling pathways that mediate CMECs dysfunction induced by hypoxia or ischemia. Akt, Protein kinase B; ALDH, Aldehyde dehydrogenase; ATP, Adenosine triphosphate; bFGF, Basic fibroblast growth factor; CCM, Compound Chinese medicine; CL, Carthamus tinctorius L. and Lepidium apetalum Willd; DGBX, DangGuiBuXue Tang; EGb761, Ginkgo biloba extract; eNOS, Endothelial nitric oxide synthase; ERK, Extracellular signal-regulated kinase; DG, GeGen DanShen extract; HIF, Hypoxia-inducible factor; IRF5, Interferon regulatory factor 5; KLF, Krüppel-like factor; MDG-1, A water-soluble beta-D-fructan from O. japonicus; MI, Myocardial infarction; mTOR, Mammalian target of rapamycin; NRG-1, Neuregulin-1; OGD, Oxygen glucose deprivation; PI3K, Phosphoinositide 3-kinase; QLQX, QiLiQiangXin; QSYQ, QiShenYiQi; S1P, Sphingosine 1 phosphate; STDP, Shexiang Tongxin Dropping Pill; SXBX, SheXiangBaoXin Pill; Syk, Spleen tyrosine kinase; TGF, Transforming growth factor; TJs, Tight junctions; TXL, TongXinLuo; VA, Velvet Antler; VAP, Velvet Antler Proteins; VEGF, Vascular endothelial growth factor; VEGFR, Vascular endothelial growth factor receptor.

**Table 1.**
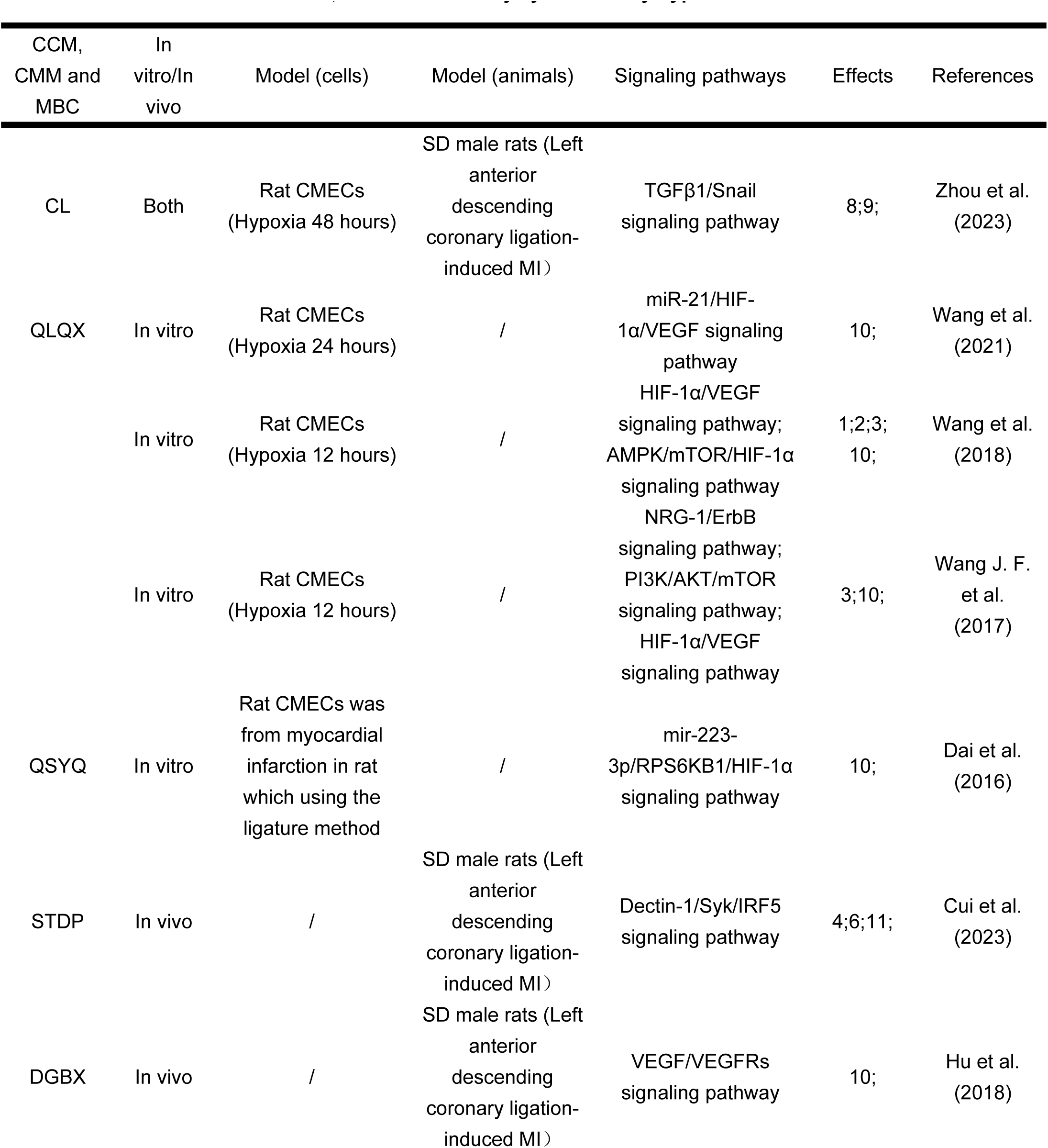
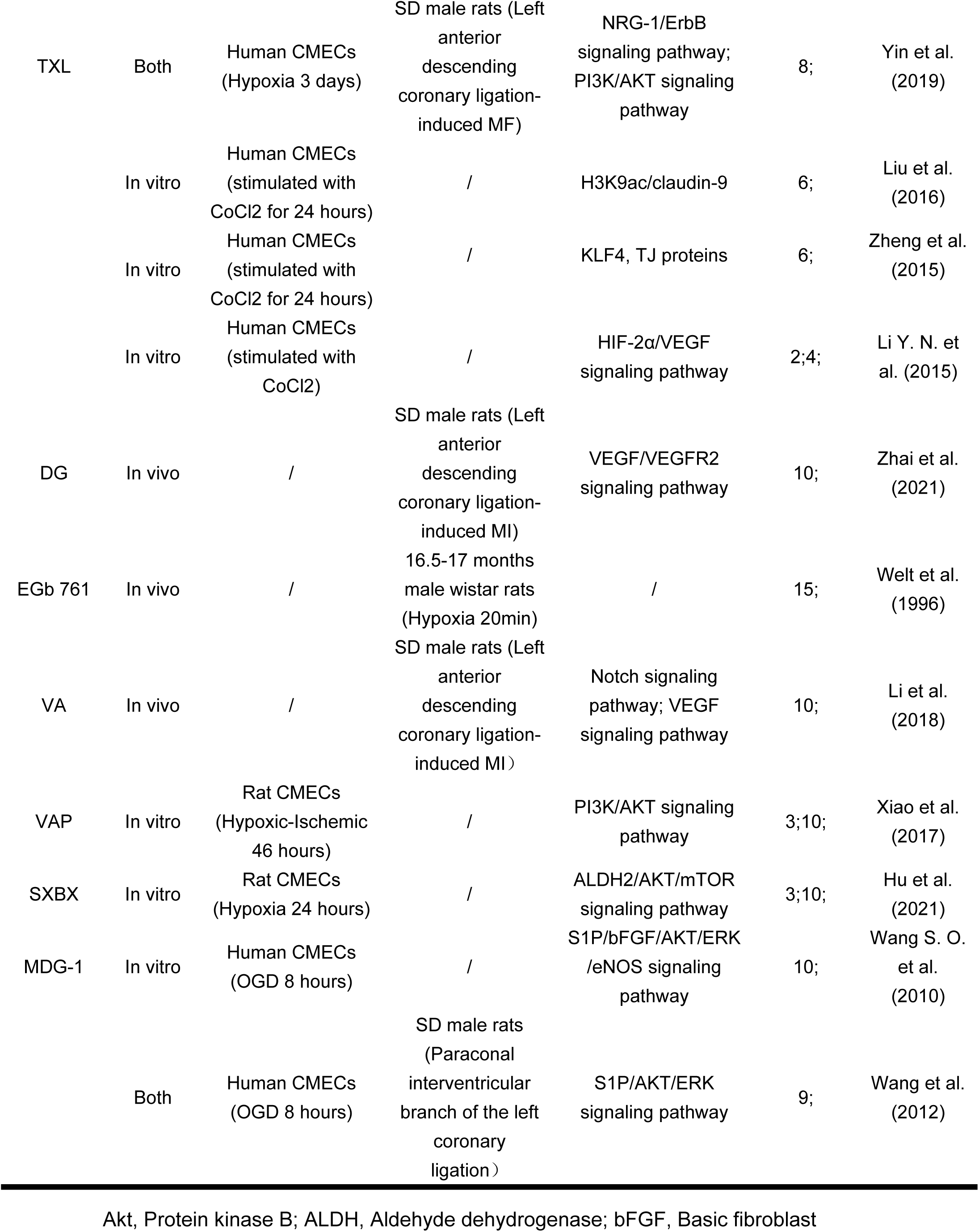
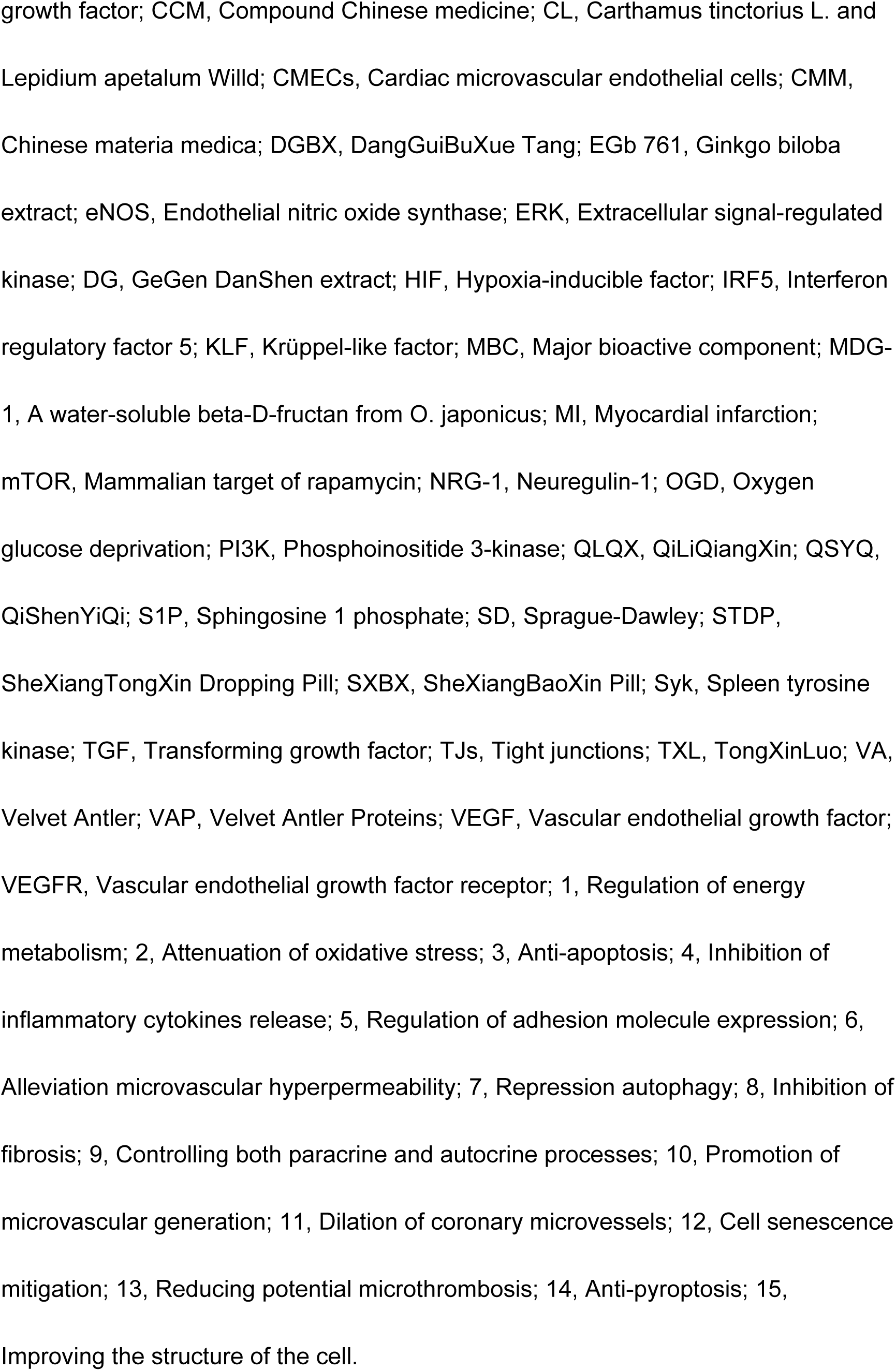
Effects of CCM, CMM and MBC injury induced by hypoxia or ischemic.

Ischemia or hypoxia can potentially impede angiogenesis. Animal studies have shown that Velvet Antler (VA), GeGen DanShen extract (DG), and DangGuiBuXue Tang (DGBX) can effectively mitigate the damage caused by myocardial infarction (MI) and enhance microvessel density within the infarct zone [17–19]. SheXiangTongXin Dropping Pill (STDP) demonstrated the ability to prevent microvascular leakage, reduce edema, hemorrhage, and inflammatory cell infiltration in the tissues surrounding the microvessels, while preserving the morphological integrity of myocardial microvessels [20]. Another study discovered that Ginkgo biloba extract (EGb761) can protect the ultrastructure of CMECs against hypoxia [21].

Hypoxia-inducible factor (HIF) is recognized as the major transcriptional regulator of the adaptive response to hypoxia [22]. The presence of HIF-1α stimulates the expression of various angiogenic factors, including transforming growth factor (TGF)-α, platelet-derived growth factor (PDGF), and vascular endothelial growth factor (VEGF) [23]. QiLiQiangXin (QLQX) has been shown to protect CMECs from hypoxia-induced injury by promoting angiogenesis and CMECs proliferation through the activation of the miR-21/HIF-1α/VEGF axis and HIF-1α-mediated glycolysis promotion [24]. In an alternative study, QLQX was found to induce the adenosine 5’-monophosphate-activated protein kinase (AMPK)/mammalian target of rapamycin (mTOR)/HIF-1α signaling pathway, leading to increased HIF-1α protein expression in hypoxic CMECs. The stability of HIF-1α and its activation of genes are closely associated with prolyl hydroxylase domain enzymes (PHDs) [25], and QLQX was found to downregulate the expression of PHDs, thereby enhancing HIF-1α stability [26]. Neuregulin-1 (NRG-1), a cardioactive growth factor released from endothelial cells, is indispensable for cardiac development, structural maintenance, and functional integrity of the heart. They transmit their signals through interactions with cell membrane receptors of the ErbB family [27]. QLQX activates the phosphatidylinositol 3-kinase (PI3K)/protein kinase B (AKT)/mTOR signaling pathway, which further activates the HIF-1α/VEGF signaling pathway to protect against hypoxia. This mechanism is mediated by the NRG-1/ErbB signaling pathway [28].

TongXinLuo (TXL) has been previously investigated as a potential treatment for luo illness. Both cyclooxygenase-2 (COX-2) and inducible nitric oxide synthase (iNOS) are inducible enzymes involved in oxidative stress and inflammation [29, 30]. In hypoxia-induced human CMECs, TXL was found to inhibit the production of VEGF, HIF-2α, COX-2, and iNOS. High doses of TXL were found to inhibit hypoxia-induced increases in the levels of the inflammatory mediator prostaglandin E2 (PGE2) and the oxidative marker nitrotyrosine (NT), thereby attenuating inflammation and oxidative damage. However, TXL did not enhance the expression of prostacyclin synthase (PGIS) or endothelial nitric oxide synthase (eNOS) [31]. PGIS possesses anti-inflammatory properties and cytoprotective effects [32], while eNOS expression promotes blood vessel dilation [33]. Macrophages have been implicated in hypoxia-induced injury to human CMECs, and peroxynitrite is involved in this process. Another study found that TXL can increase the expression of PGIS primarily and reduce endothelin-converting enzyme-1 (ECE-1) expression through inhibiting macrophage-mediated nitrotyrosine accumulation [34]. Notably, ECE-1 plays a crucial role in the signaling pathway for endothelin and serves as a significant modulator of vascular tone [35].

The integrity of the microvascular barrier relies on various factors, including the subendothelial basement membrane, caveolin quantity and function in endothelial cells, and the presence of adherens junctions (AJs) and tight junctions (TJs) between adjacent CMECs [36]. TJs proteins encompass occludin, claudin, and junctional adhesion molecule (JAM), which are transmembrane proteins [37] AJs are responsible for calcium-mediated homophilic adhesion between neighboring cells through cadherin, including zona occludens 1 (ZO-1) protein, β-catenin, vascular endothelial (VE)-cadherin, among others [38, 39]. TXL can reverse the hypoxia-inhibited claudin-9 expression by elevating histone H3 lysine 9 acetylation (H3K9ac) in the promoter region of its gene [40]. Furthermore, TXL provides protection by enhancing the phosphorylation of krüppel-like factor (KLF) 4, a basic transcription factor that regulates the production of occludin, claudin-1, VE-cadherin, and β-catenin [41].

The transcriptional repressor snail can induce the endothelial-to-mesenchymal transition (EndMT). The resultant EndMT-derived cardiac fibroblasts (CFs), synthesize large amounts of collagen, which contributes to fibrotic lesions and abnormal collagen fiber deposition [42]. TXL inhibits EndMT by downregulating snail expression in CMECs after three days of hypoxia and activates the NRG-1/ErbB/PI3K/AKT signaling pathway to reduce myocardial fibrosis (MF) following acute MI [43]. Hypoxia for 48 hours activates the TGF-β1/snail signaling pathway, and the Carthamus tinctorius L. and Lepidium apetalum Willd drug pair (CL) can inhibit EndMT in CMECs by inhibiting the TGF-β1/snail signal pathway [44].

QiShenYiQi (QSYQ) downregulate the expression of mir-223-3p, activate the ribosomal protein S6 kinase B1 (RPS6KB1)/HIF-1α signaling pathway, and facilitate ischemic cardiac angiogenesis [45]. A study on Ophiopogon japonicus and its water-soluble beta-D-fructan (MDG-1) revealed that the sphingosine 1 phosphate (S1P)/AKT/extracellular signal-regulated kinase (ERK) signaling pathway might be involved in the potential mechanism of their anti-ischemic action on cell survival [46]. Basic fibroblast growth factor (bFGF) is a prominent factor that stimulates endothelial cell proliferation and neovascularization by binding to the fibroblast growth factor (FGF) receptor [47]. Another investigation demonstrated that MDG-1 could promote angiogenesis by protecting human CMECs and cardiomyocytes from ischemia-induced cellular damage through the S1P/bFGF/AKT/ERK/eNOS signaling pathway [48]. SheXiangBaoXin Pill (SXBX) promotes angiogenesis through the aldehyde dehydrogenase (ALDH)2/AKT/mTOR signaling pathway [49]. Velvet antler proteins (VAP) safeguards CMECs against ischemia-hypoxia by modulating the PI3K/AKT signaling pathway [50].

### 3.2. TCM that Modify CMECs in I/R or H/R Injury Model

I/R-induced damage to the microcirculation can lead to increased leukocyte adhesion, reactive oxygen species (ROS) production, dysfunction of CMECs, and leakage of macromolecules. These events can trigger inflammatory responses, impair CBF, and ultimately result in cardiomyocyte death. Therefore, the preservation of CMECs and the reduction of CMD following I/R are considered promising approaches in the treatment of cardiac I/R injury [51].

Studies have demonstrated that TCM can inhibit apoptosis, regulate the expression of adhesion molecules, alleviate microvascular permeability, dilate coronary microvessels, mitigate microvascular obstruction, modulate oxidative stress, and control inflammation, among other effects. According to research on animals, the DanLou formula (DLF) and the TongMaiYangXin pill (TMYX) enhance myocardial no-reflow following I/R [52, 53]. Resveratrol (RSV) and scutellarin (SCU) have been found to regulate the production of intracellular proteins and the paracrine secretion function of CMECs following H/R damage [54, 55]. The specific effects and underlying mechanisms of action are illustrated in Fig 3 and summarized in Table 2.

**Fig 3.**
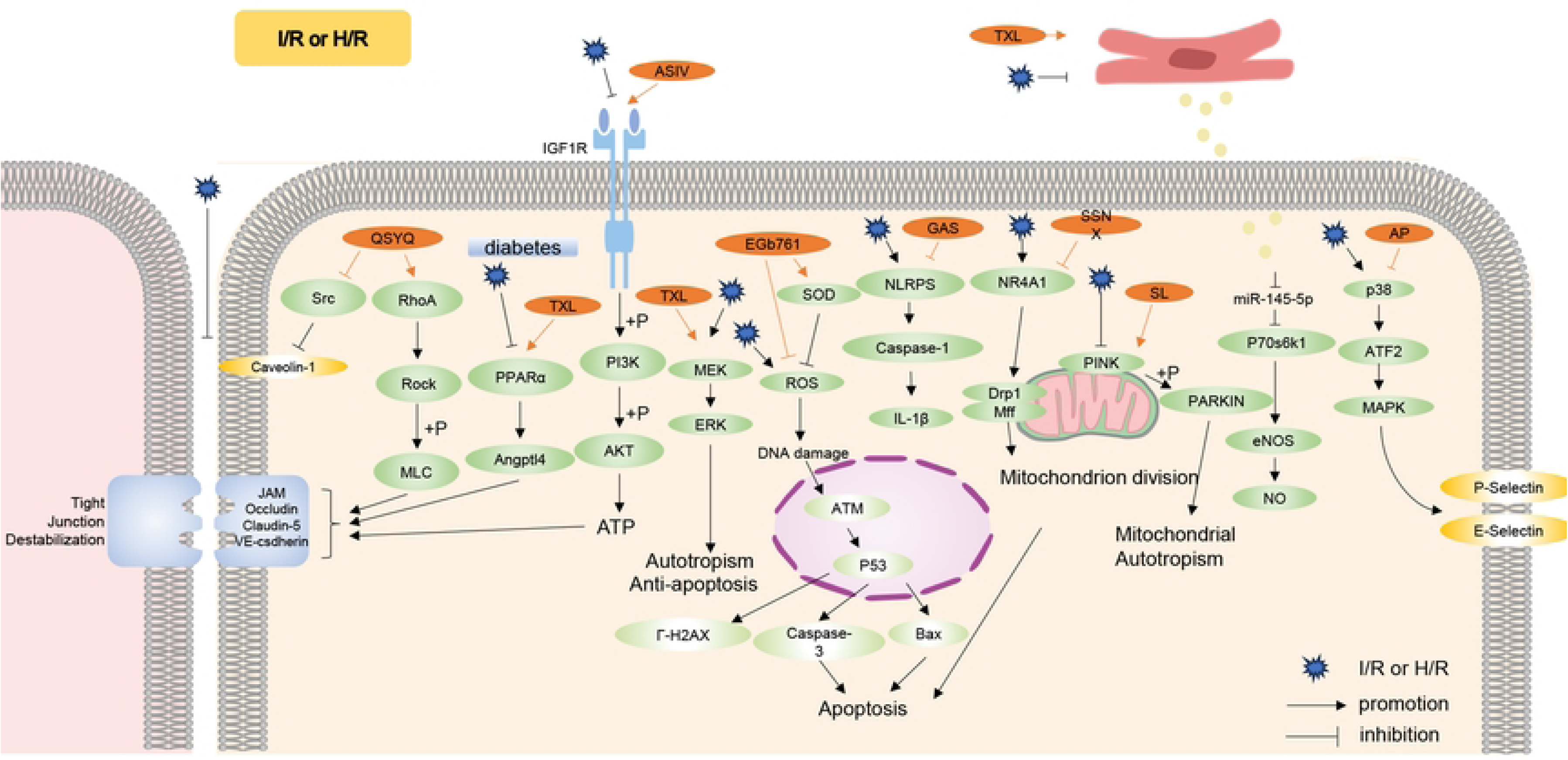
TCM regulate various signaling pathways that mediate CMECs dysfunction induced by I/R or H/R. AKT, Protein kinase B; AP, Astragalus polysaccharide; ASIV, Astragaloside IV; ATM, Ataxia telangiectasia mutated; ATP, Adenosine triphosphate; Drp1, Dynamin-related protein 1; EGb761, Ginkgo biloba extract; eNOS, Endothelial nitric oxide synthase; ERK, Extracellular signal-regulated kinase; GAS, Gastrodin; H/R, Hypoxia/Reoxygenation; I/R, Ischemia/reperfusion; IGF1R, Insulin-like growth factor 1 receptor; iNOS, Inducible nitric oxide synthase; IRS1, Insulin receptor substrate 1; MAPK, Mitogen-activated protein kinase; MEK, Mitogen-activated protein kinase; Mff, Mitochondrial fission factor; NLRP3, Nucleotide-binding oligomerization domain-like receptor protein 3; NR4A1, Nuclear receptor subfamily 4 group A member 1; PI3K, Phosphoinositide 3-kinase; PINK, Phosphatase and tensin homolog-induced putative kinase; QSYQ, QiShenYiQi; SCU, Scutellarin; SL, ShenLian extract; SSNX, ShuangShenNingXin formula; TXL, TongXinLuo.

**Table 2.**
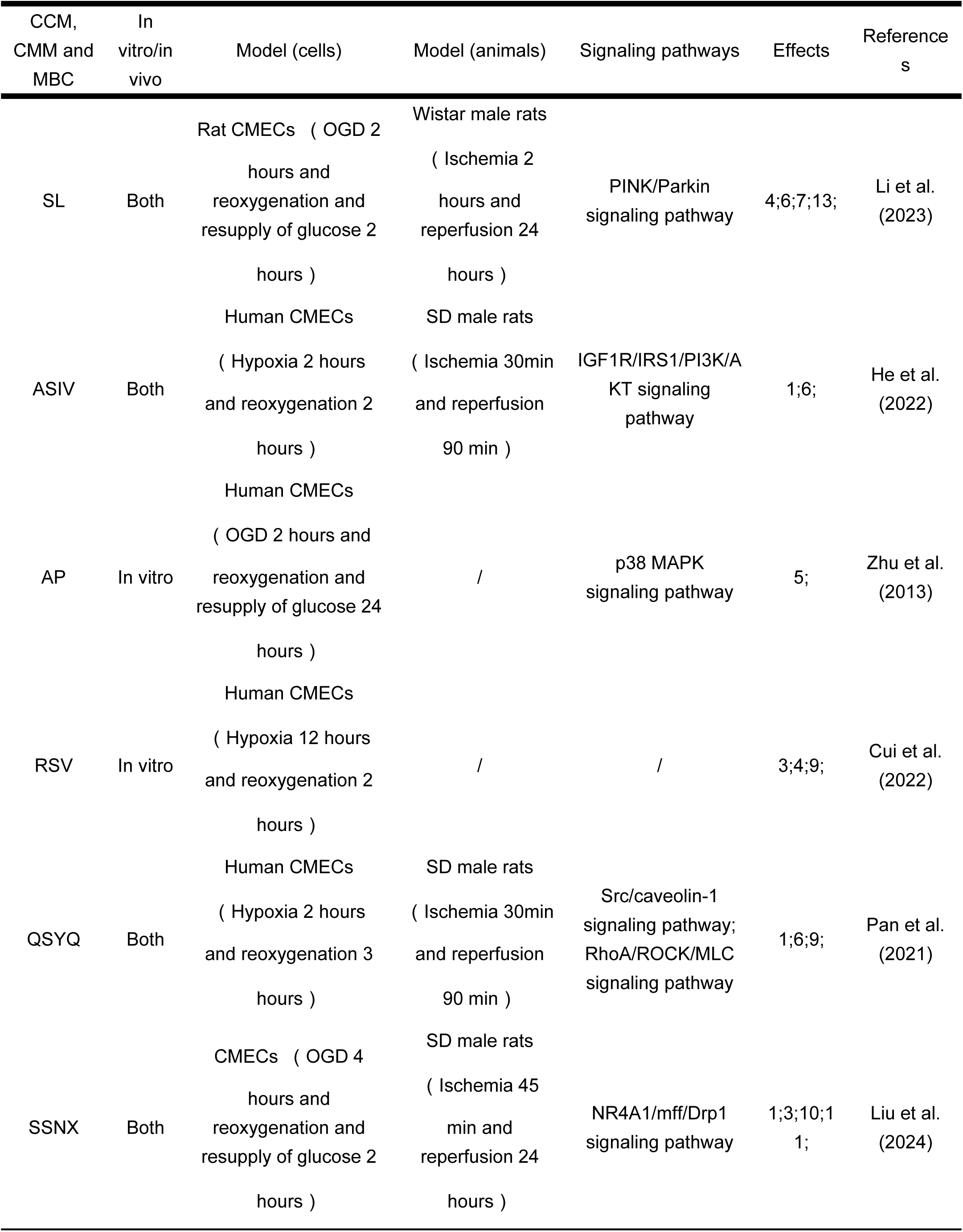
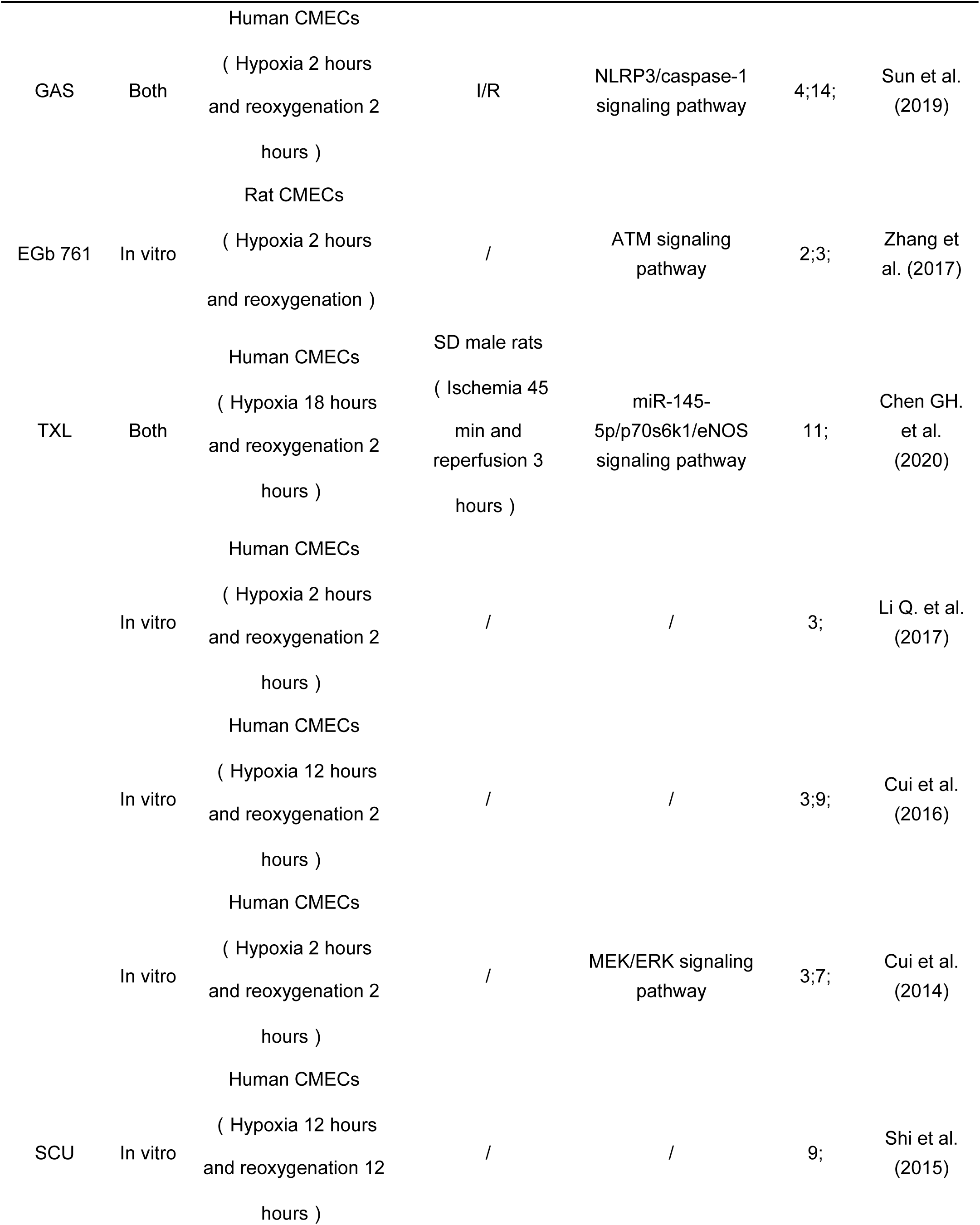
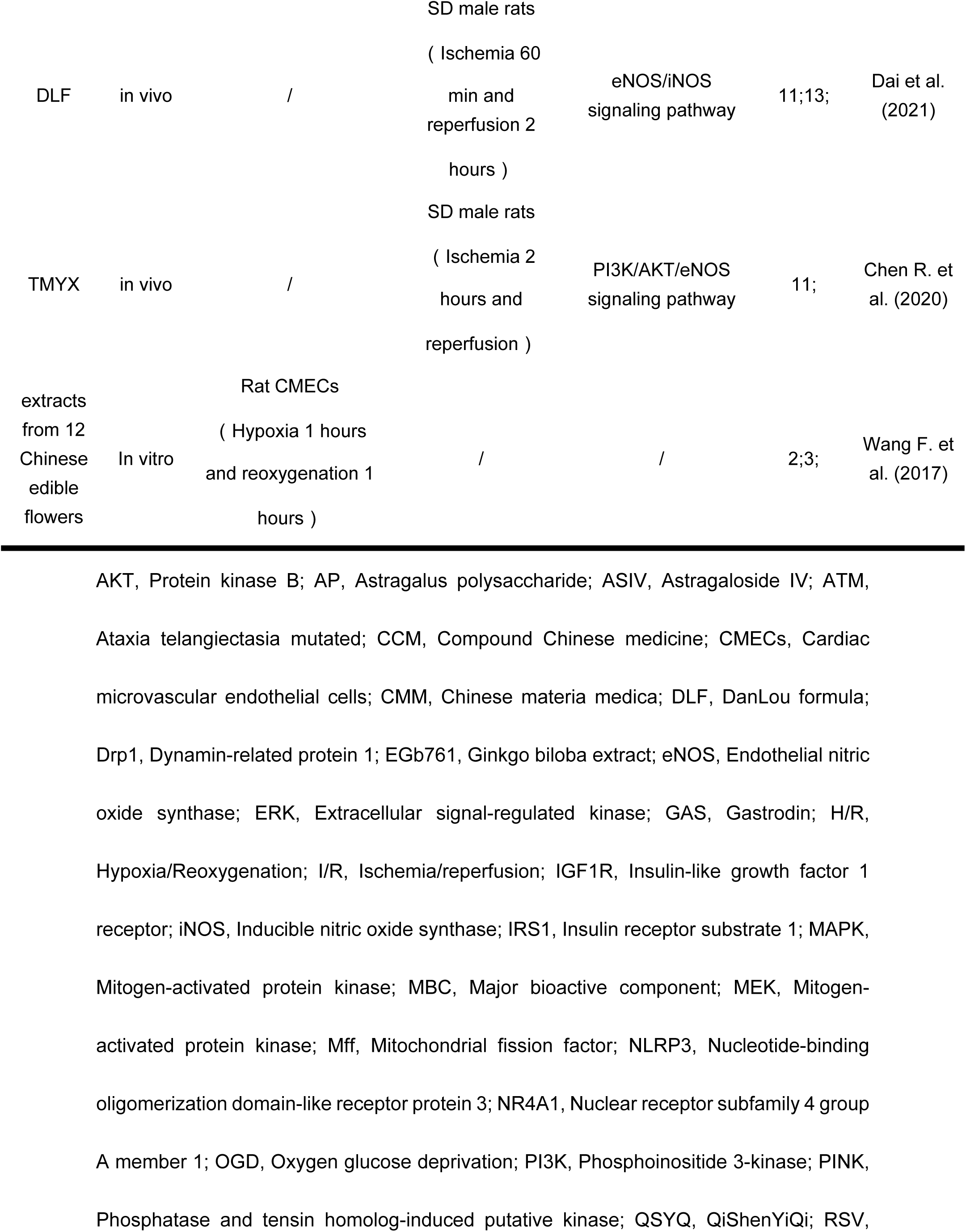
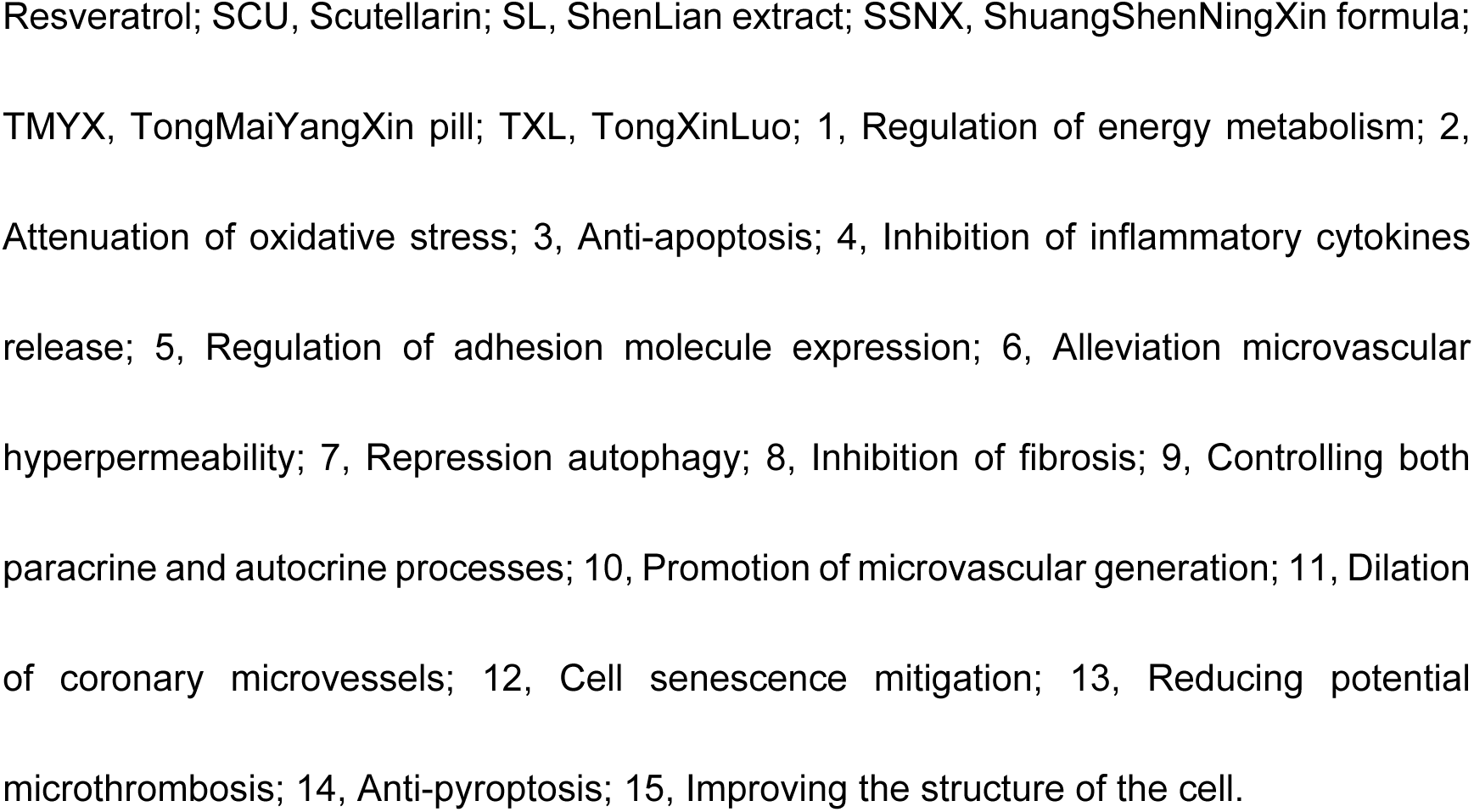
Effects of CCM, CMM and MBC injury induced by I/R or H/R.

Through the ras homolog gene family member A (RhoA)/Rho-associated protein kinase (ROCK)/myosin light chain (MLC) signaling pathway, QSYQ have been found to inhibit I/R-induced cardiac microvascular hyperpermeability. Src family protein tyrosine kinases are known to be associated with endothelial cell permeability [56]. QSYQ has been shown to attenuate the upregulation of Src, p-caveolin-1, matrix metallopeptidase-9 (MMP-9), and cathepsin S (CTSS) induced by H/R, while also preserving the expression of claudin-5. Among the signaling pathways involved in mediating the effects of QSYQ, the Src/caveolin-1 pathway has been implicated [57].

Astragaloside IV (ASIV), the main active ingredient in QSYQ, has been found to protect the microvascular endothelial barrier. ASIV mitigates adenosine triphosphate (ATP) depletion, enhances the expression of tight junction proteins between endothelial cells, and promotes the H/R-induced activation of the insulin-like growth factor 1 receptor (IGF1R) and downstream phosphorylation of insulin receptor substrate 1 (IRS1)/PI3K/AKT signaling pathway [58].

Following I/R, the release of cytokines, oxygen radicals, and pro-inflammatory mediators stimulates the vascular endothelium and neutrophils. This leads to the upregulation of adhesion molecules, promoting the adhesion and migration of white blood cells across the vascular endothelium [59]. Astragalus polysaccharide (AP), a key component of QSYQ, reduces the expression of relevant adhesion molecules following I/R. AP also inhibits the interaction between human CMECs and polymorphonuclear leukocyte (PMN) during I/R by suppressing the p38 mitogen-activated protein kinase (MAPK) signaling pathway and downregulating the expression of adhesion molecules (P-selectin and E-selectin) in human CMECs [60].

TXL has been shown to exert a protective effect against H/R injury, inhibit apoptosis in CMECs, and regulate protein expression and paracrine function in these cells [61, 62]. In cardiomyocytes treated with TXL following H/R, vesicles containing long intergenic non-coding RNA regulator of reprogramming (Linc-ROR) are released and taken up by CMECs. Subsequently, Linc-ROR downregulates its target miR-145-5p, which in turn promotes the production of 70 kDa ribosomal protein S6 kinase 1 (P70S6k1) and activates the eNOS pathway in CMECs [63]. Autophagy is an essential cellular process involved in the degradation of aging or dysfunctional organelles and protein aggregates, serving as a quality control mechanism. Impaired autophagy leads to the accumulation of dysfunctional organelles and proteins, resulting in endoplasmic reticulum stress (ERS) and apoptosis [64]. TXL has been found to induce autophagy via activation of the mitogen-activated protein kinase (MEK)/ERK signaling pathway, thereby protecting human CMECs from H/R injury [65].

I/R injury in CMECs induces signaling pathways involved in both mitochondrial division and apoptosis. ShenLian extract (SL) has been shown to prevent mitochondrial autophagy and preserve mitochondrial activity, thereby reducing endothelial cell death, preserving endothelial cell function, and protecting the microvasculature, which ultimately helps mitigate coronary artery no-reflow. The phosphatase and tensin homolog-induced putative kinase (PINK)/Parkin signaling pathway is implicated in this process [66].

ShuangShenNingXin formula (SSNX) restores mitochondrial division to normal levels and inhibits mitochondrial apoptosis, thereby reducing potential damage to the mitochondrial membrane and preventing its opening. The mechanism underlying the effects of SSNX may involve the nuclear receptor subfamily 4 group A member 1 (NR4A1)/mitochondrial fission factor (Mff)/dynamin-related protein 1 (Drp1) signaling pathway [67]. NR4A1 has been shown to regulate mitochondrial fission [68]. Mff is known to stimulate mitochondrial fission by recruiting Drp1, a critical protein involved in the control of mitochondrial division. Pathological mitochondrial fission in CMECs can trigger unfavorable mitochondrial apoptotic pathways [69].

Oxidative stress is a crucial factor in I/R injury, and ROS play a central role in mediating oxidative stress [70]. EGb761 has been shown to inhibit the I/R-induced activation of the ataxia telangiectasia mutated (ATM) pathway, thereby ameliorating apoptosis by suppressing ROS expression [71]. In a study evaluating the antioxidant effects of 12 edible flowers. Most of the edible flowers examined demonstrated the potential to enhance the antioxidant capacity of CMECs following I/R. Honeysuckle, rose, and wild chrysanthemum were found to exhibit the highest levels of antioxidant activity [72].

The nucleotide-binding oligomerization domain-like receptor protein 3 (NLRP3)/caspase-1 signaling pathway is believed to play a critical role in the injury to CMECs and cardiac tissues, as well as the increase in infarct area during I/R injury. Gastrodin (GAS) has been shown to partially reverse the pyroptosis of CMECs and reduce the area of MI and inflammatory cell infiltration by inhibiting the NLRP3/caspase-1 signaling pathway [73].

### 3.3. TCM that Modify CMECs in Inflammatory Injury Model

Vascular inflammation plays a significant role in the pathogenesis of CMD [74], It can lead to damage in coronary microvessels, as well as promote thrombosis and perivascular fibrosis. The vascular endothelium, acting as a barrier between the vascular lumen and surrounding tissue, serves as a key regulator and participant in the vascular inflammatory response [75]. Inflammatory substances such as tumor necrosis factor-α (TNF-α), homocysteine (Hcy), and lipopolysaccharide (LPS) are known to induce inflammation in cells [76–78]. TCM exerts its effects on enhancing CMECs following vascular inflammatory injury mainly through modulation of the nuclear factor-kappa B (NF-κB), MAPKs, and Janus tyrosine kinase (JAK)/signal transducer and activator of transcription (STAT) signaling pathways [79, 80]. For instance, ShenMai formula (SMF), 4-O-(2-O-acetyl-6-O-p-coumaroyl-β-D-glucopyranosyl)-p-coumaric acid (4-ACGC) isolated from Bidens pilosa Linn., and salidroside (SA) have been shown to impact the development of vascular inflammation [76, 81, 82]. Fig 4 and Table 3 provide a visual representation of the underlying mechanisms and the expression of relevant inflammatory mediators.

**Fig 4.**
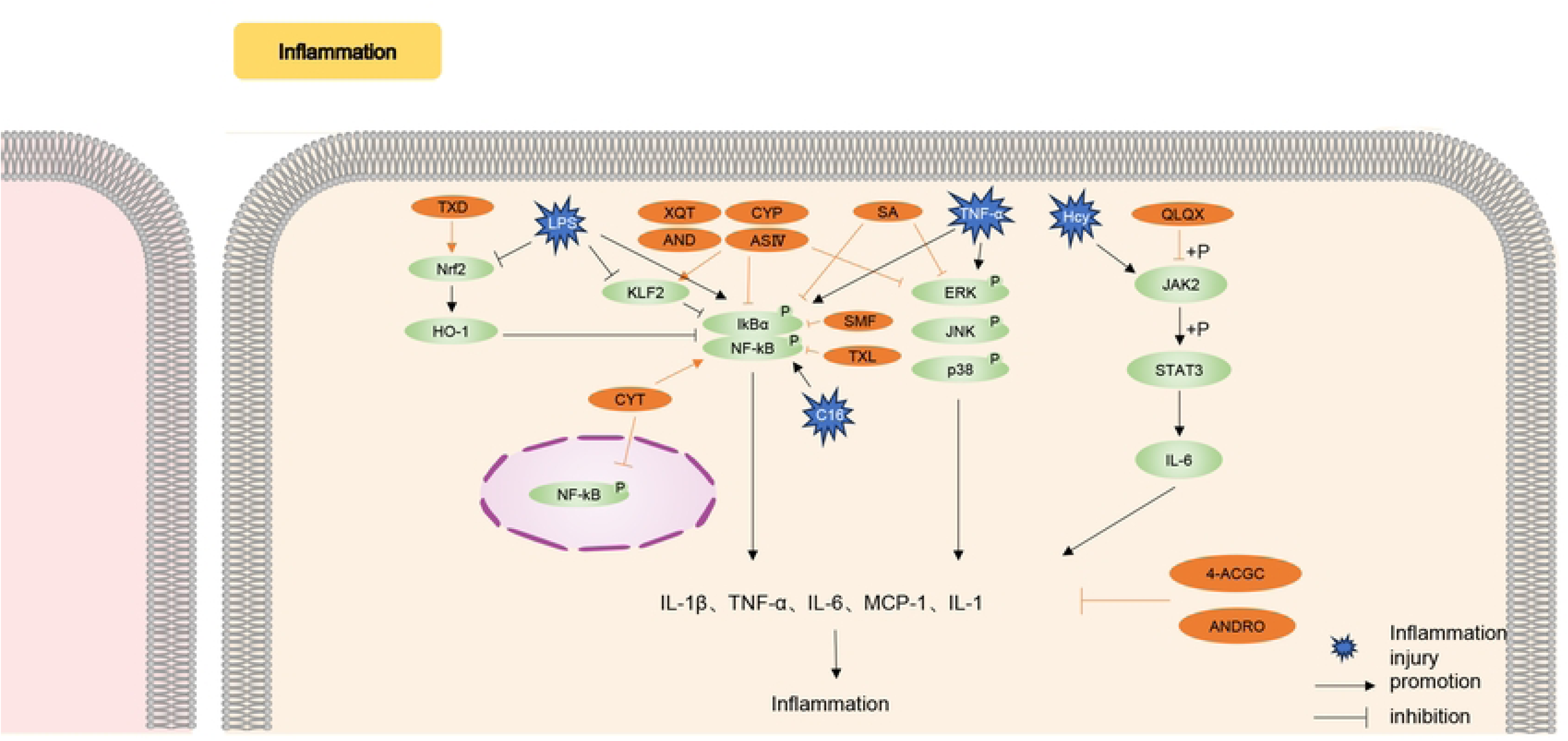
TCM regulate various signaling pathways that mediate CMECs dysfunction induced by inflammation. 4-ACGC, 4-O-(2″-O-acetyl-6″-O-p-coumaroyl-β-D-glucopyranosyl) -p-coumaric acid; AND, Andrographolide; ASIV, Astragaloside IV; CYP, Alpha-Cyperone; CYT, Caffeoylxanthiazonoside; Hcy, Homocysteine; HO-1, Heme oxygenase-1; IL, Interleukin; JAK, Janus tyrosine kinase; KLF, Krüppel-like factor; LPS, lipopolysaccharide; MAPK, Mitogen-activated protein kinase; MDA, Malondialdehyde; NF-κB, Nuclear factor-kappa B; Nrf2, Nuclear factor-erythroid 2 related factor 2; QLQX, QiLiQiangXin; SA, Salidroside; SD, Sprague Dawley; SMF, ShenMai formula; SOD, Superoxide Dismutase; STAT, Signal transduction and activator of transcription; TNF-α, Tumor necrosis factor-α; TNF-α, Tumor necrosis factor-α; TXD, TianXiangDan; TXL, TongXinLuo; XQT, XiangQiTang.

**Table 3.**
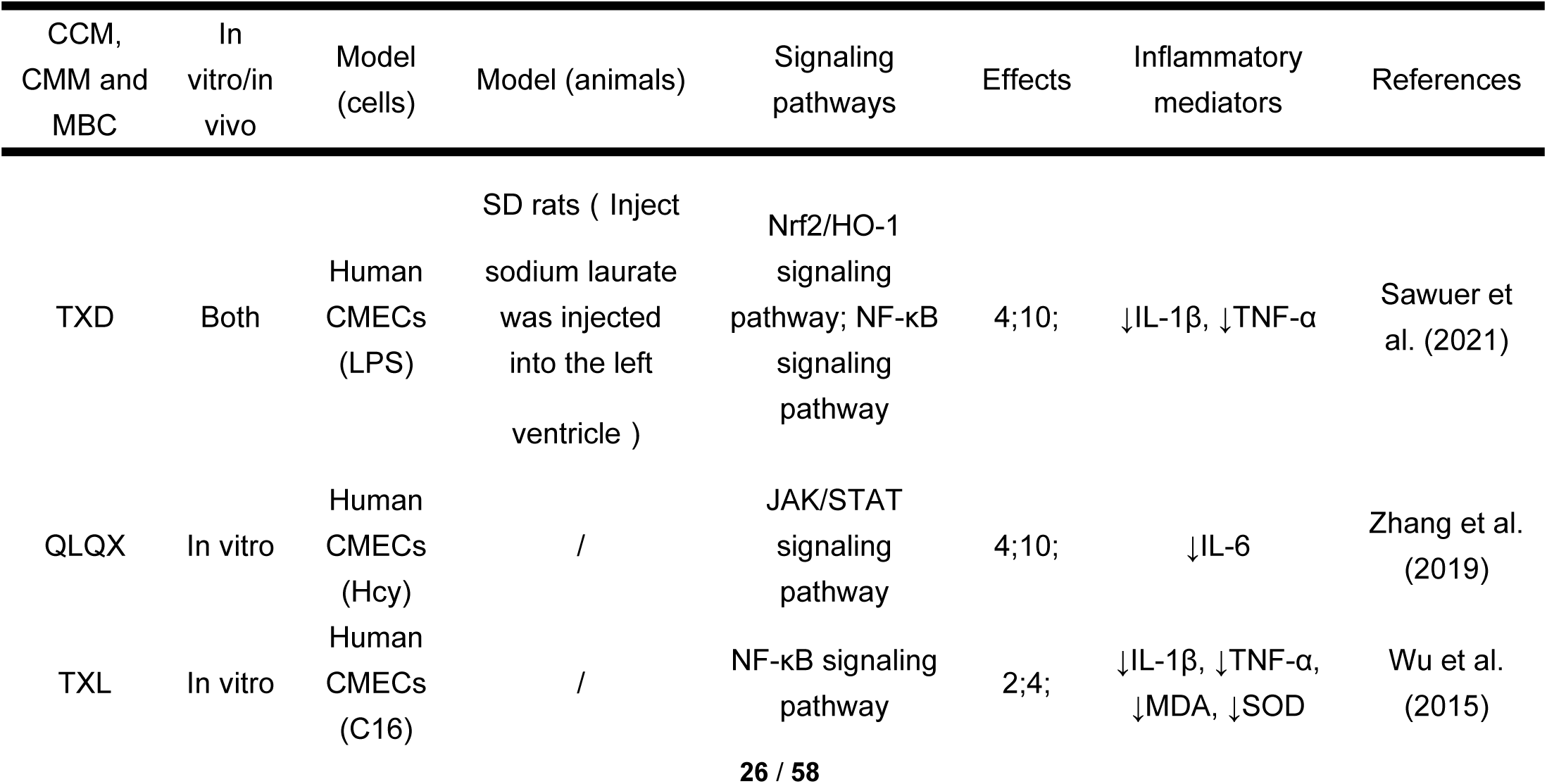
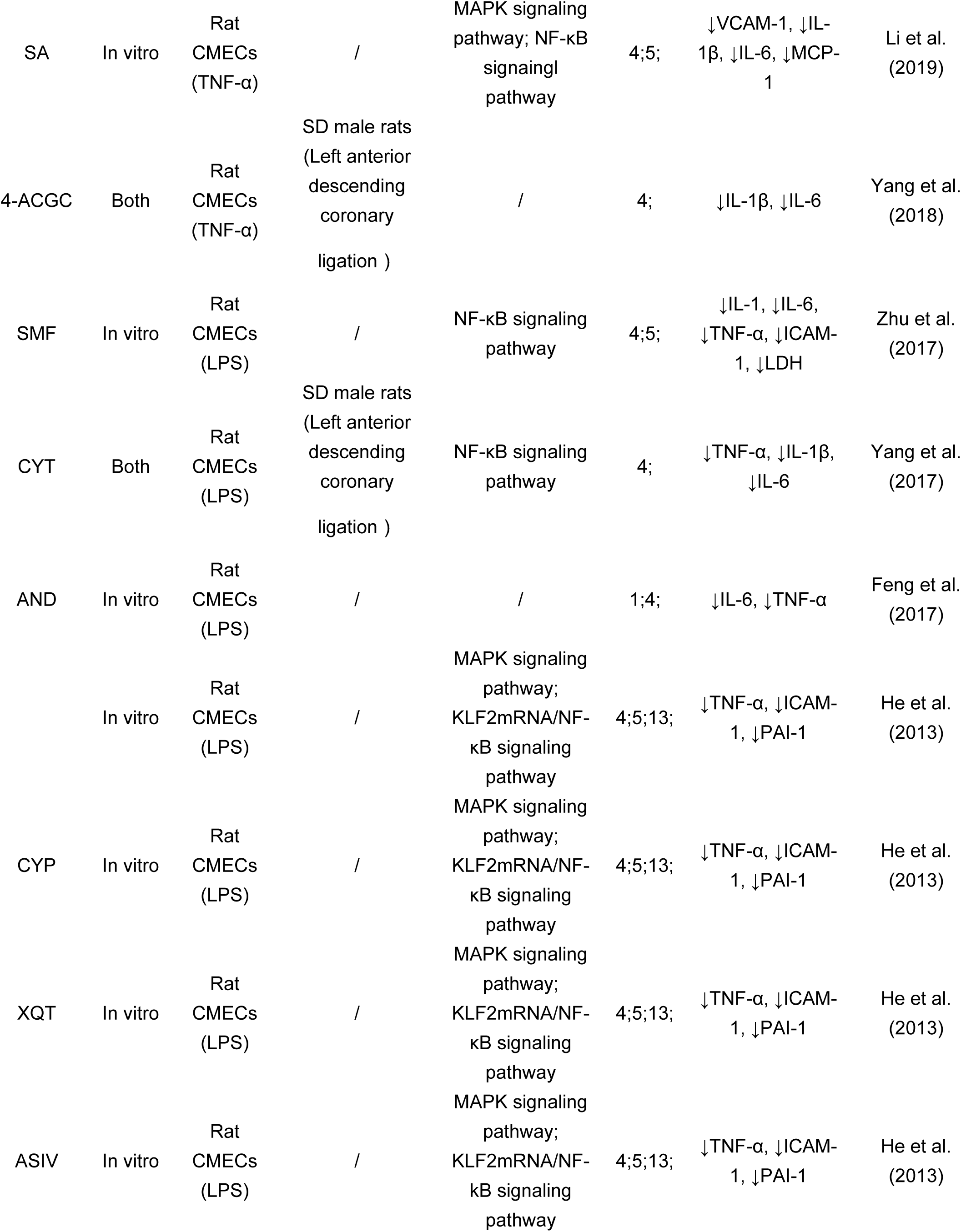
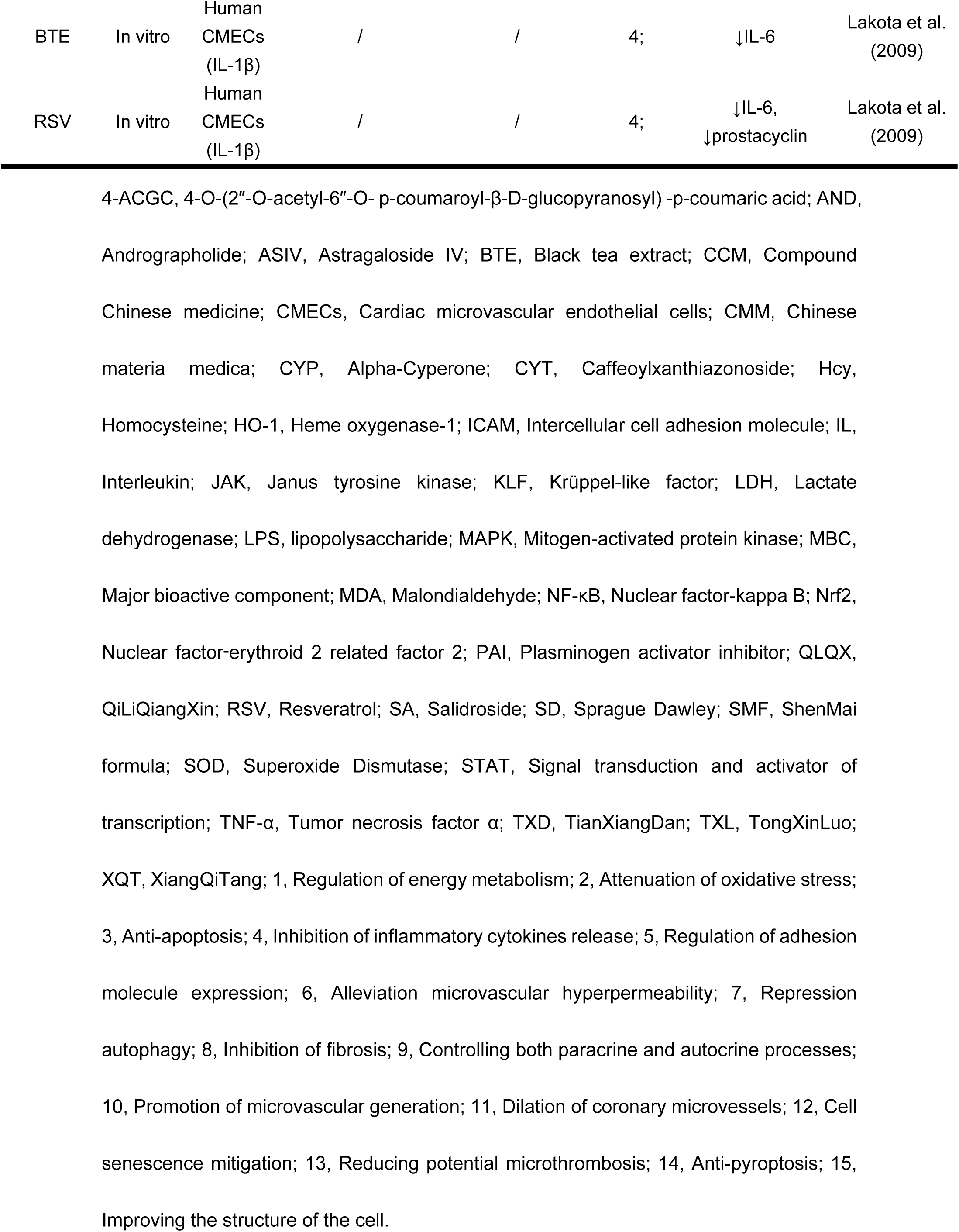
Effects of CCM, CMM and MBC injury induced by inflammatory.

The inhibitory effect of TianXiangDan (TXD) on LPS-induced microvascular endothelial inflammation has been associated with the activation of nuclear factor-erythroid 2-related factor 2 (Nrf2). Heme oxygenase-1 (HO-1) is one of the downstream proteins in the Nrf2 signaling pathway. TXD exerts its anti-inflammatory effects by suppressing the expression of TNF-α, phosphorylated inhibitor of kappa B alpha (p-IκBα), phosphorylated p65 (p-p65), and interleukin (IL)-1β through the induction of HO-1 protein [83].

C16 induction in cells leads to oxidative stress and inflammation. Among the various sources of intracellular ROS, Nicotinamide adenine dinucleotide phosphate (NADPH) oxidase plays a significant role [84, 85]. TXL inhibits the release of TNF-α and IL-1β induced by C16 through the blockade of NF-κB activation and expression. Moreover, TXL suppresses the upregulation of malondialdehyde (MDA) and superoxide dismutase (SOD), thereby reducing the production of ROS. Its antioxidant activities may be attributed to the suppression of HO-1 and NADPH oxidase complex expression in human CMECs [86].

Different levels of NF-κB p65 expression and regulation are observed in the cytoplasm and nucleus. When exposed to LPS, the fruit of Xanthium strumarium L plant contains an active ingredient called caffeoylxanthiazonoside (CYT), which significantly reduces the production of TNF-α, IL-1β, and IL-6. Additionally, there is an upregulation of inhibitor of NF-κB (IκB) and cytoplasmic NF-κB p65 protein expression, along with a downregulation of nuclear NF-κB p65 [87].

Inflammation and coagulation are closely related and can mutually promote each other [88]. Plasminogen activator inhibitor (PAI)-1 [89] and tissue factor (TF) [90] are important factors in the coagulation cascade. XiangQiTang (XQT) and its components, Alpha-Cyperone (CYP), ASIV, and andrographolide (AND), inhibit LPS-stimulated production of TNF-α, intercellular adhesion molecule-1, and PAI-1. They also upregulate KLF2 mRNA expression, reduce the phosphorylation level of NF-κB p65 protein, and inhibit TF secretion. Moreover, XQT, CYP, ASIV, and AND suppress the expression of proteins involved in the MAPK signaling pathway [91].

Caveolin-1 is a crucial structural protein that facilitates the transmembrane transport of low-density lipoprotein cholesterol (LDL-C), thereby promoting the development of atherosclerosis [92]. AND significantly reduces caveolin-1 expression in LPS-induced CMECs. It modulates the expression of IL-6 and TNF-α among inflammatory factors and inhibits extracellular ATP-induced calcium release by decreasing the expression of phospholipase Cδ3 (PLCδ3), without affecting extracellular calcium endocytosis. This has a limited impact on nitric oxide (NO) production and release [93].

Lakota et al.[94] demonstrated that human CMECs responded dose-dependently to IL-1β-induced IL-6 levels and prostacyclin release. The administration of black tea extract (BTE) and RSV inhibited IL-1β-induced responses. Additionally, QLQX downregulated the expression of phosphorylated STAT3, phosphorylated JAK2, and IL-6 in CMECs. It upregulated the expression of vascular-endothelial growth factor A (VEGFA), mitigated the inflammatory process induced by Hcy, and promoted angiogenesis, potentially through modulation of the JAK/STAT signaling pathway [95].

### 3.4. TCM that Modify CMECs in Metabolic Injury Model

Cellular metabolism is essential for maintaining normal cellular biochemical processes and biological activities. Patients with metabolic abnormalities, such as elevated levels of free fatty acids and chronic hyperglycemia, face an augmented risk of cardiovascular events [96].

A high fat diet (HFD) is a significant risk factor for organ damage, including the liver, kidneys, and heart, leading to increased mortality [97]. HFD can induce capillary permeability in the microvascular environment, potentially leading to interstitial fibrosis and myocardial dysfunction [98]. In mice fed a HFD, chronic intermittent administration of quercetin (Q) reduces intramyocardial fat accumulation, increases cardiac microvessel density, and regulates oxidative stress [99].

Apolipoprotein E (apoE) deficiency is known to cause elevated levels of cholesterol-rich compounds in the bloodstream, contributing to the development of atherosclerotic lesions [100]. Treatment with TXL significantly decreases lipid levels, reduces atherosclerotic plaque formation in apoE-deficient mice, and improves endothelial cell function [86].

In animal models, N(ω)-nitro-L-arginine methyl ester (L-NAME), a nitric oxide synthase (NOS) inhibitor, suppresses NOS activity and reduces NO production, leading to increased blood pressure and the progression of left ventricular remodeling [101]. QSYQ has been found to increase myocardial capillary density and inhibit microvascular endothelial inflammation induced by L-NAME combined with the HFD [102].

A high glucose (HG) environment can downregulate the expression of claudins-5 and -11 in human CMECs. TXL can reverse the HG-induced inhibition of claudins-5 and -11 by increasing H3K9ac in the promoters of these genes. Moreover, high-dose TXL treatment promotes the membrane localization of claudins-5 and -11 in HG-stimulated human CMECs [103].

Angiopoietin-like 4 (Angptl4) plays a protective role in regulating the endothelial barrier, maintaining vascular integrity by preserving the VE-calmodulin complex [104]. In the context of preserving the structure and function of the endothelial barrier under conditions of high glucose-induced I/R, TXL has been found to be comparable to insulin and recombinant human Angptl4. The expression of Angptl4 can be induced by peroxisome proliferator-activated receptor α (PPAR-α). In diabetic patients, TXL may preserve the integrity of the endothelial barrier against I/R injury through the activation of the PPAR-α/Angptl4 signaling pathway [105, 106].

### 3.5. TCM that Modify CMECs in Ang II Injury Model

Ang II is a bioactive peptide that regulates vascular tone and promotes the proliferation of vascular smooth muscle cells, playing a pivotal role in the pathogenesis of cardiovascular disease. Ang II has been implicated in inducing cardiac hypertrophy [107], a which is an independent risk factor for mortality [108]. Studies have found that Ang II-induced apoptosis in CMECs is closely associated with the development of CMD in heart failure patients [109]. It has been observed that Ang II enhances endothelial cell apoptosis, impairs the shear response of CMECs, and hampers the morphological adaptation to shear stress by downregulating the expression of platelet-endothelial cell adhesion molecule-1 (PECAM-1). Allicin (A), through stimulation of the PECAM-1/PI3K/AKT/eNOS signaling pathway, downregulation of caspase-3 and receptor interacting protein 3 (RIP3) expression, and prevention of necrotic apoptosis, enhances the functionality of CMECs. A was found to increase microvessel density in rats with cardiac hypertrophy induced by abdominal aortic constriction [110].

Autophagy is a cyclic process involved in maintaining cellular homeostasis [111]. The Forkhead-Box Class O (FoxO) family member, FoxO3a, regulates autophagy by activating genes involved in autophagosome formation [112]. CMECs exposed to Ang II undergo apoptosis, but QLQX prevents this by inhibiting autophagy through the ErbB2/AKT/FoxO3a signaling pathway [113]. In terms of the microvascular endothelial barrier, TXL attenuates the damage to human CMECs induced by Ang II by promoting KLF5 expression, which enhances the levels of tight junction proteins [114].

### 3.6. TCM that Modify CMECs in Other Injury Models

Aging is considered an independent factor associated with endothelial cell dysfunction [115]. TCM has shown promise in alleviating senescence in CMECs. From a cytoskeletal perspective, age affects the structure and function of F-actin. Extracts of Panax Notoginseng (PN), Radix Ginseng (RG), and Rhizoma Ligustici Chuanxiong (RLC) have been found to delay the senescence of CMECs in response to heat shock protein 27 (HSP27) and reduce F-actin synthesis [116].

Viral myocarditis (VMC) is a common cardiovascular disease [117]. Chronic phase VMC is characterized by MF [118]. CFs are the most affected cells in MF [119]. It has been discovered that CFs can also arise from the EndMT process, which may contribute to MF development in VMC [120]. Ginsenoside-Rb3 (Rb3), a major component of Sanqi and Renshen, inhibits EndMT in CMECs after coxsackievirus B3 (CVB3) infection through the proline-rich tyrosine kinase (Pyk) 2/PI3K/AKT signaling pathway [121]. Ginsenoside-Rg3 (Rg3) activates AKT to upregulate the Nrf2/antioxidant response element (ARE) pathway, thereby mitigating cardiotoxicity caused by adriamycin (ADM) and ameliorating endothelial dysfunction resulting from oxidative stress [122]. Millettia pulchra Kurz var.laxior (Dunn) Z. Wei is a wild plant from the Fabaceae family with diverse therapeutic uses. Its root contains the flavonoid monomer 17-Methoxyl-7-hydroxy-benzene-furanchalcone (MHBFC) [123]. In an animal model, MHBFC attenuated L-NAME-induced apoptosis of CMECs during cardiac remodeling in rats [124]. Diosmetin-7-O-β-D-glucopyranoside (Diosmetin-7-O-glucoside) is a natural flavonoid abundant in citrus fruits and herbal extracts like fructus trichosanthes peel [125]. In primary CMECs, TGF-β1 promoted EndMT. Diosmetin-7-O-glucoside, partly through an Src-dependent mechanism, regulates EndMT via endoplasmic reticulum stress [126].

Some studies have focused on the direct intervention of Chinese medicine or active ingredients without specifically addressing CMEC injury. For instance, the chemical constituents of Ophiopogon japonicus (OJ) fiber root and DG were found to modulate angiogenesis in human CMECs, promoting microvessel formation [127, 128]. In terms of regulating microvascular function, long-term oral treatment with Oroxylin A (OA), the primary constituent of Radix Scutellariae (RS), was found to enhance the production of NO and the expression of eNOS protein in CMECs, as well as the production of NO and the expression of iNOS protein in vascular smooth muscle cells (VSMCs). It was proposed that the mechanism of action of OA involved the modulation of the estrogen receptor (ER) signaling pathway [129]. Tanshinone II A (Tan II A) was found to activate the ER signaling pathway in primary CMECs, leading to increased expression of eNOS gene, NO production, ERK1/2 phosphorylation, and Ca2+ mobilization [130]. In the treatment of heart failure (HF), periplocin (PER) in Cortex Periplocae Sepii Radicis was compared with the cardiac glycoside ouabain. PER was found to increase cell proliferation, reduce cell damage, inhibit apoptosis, and affect the expression of guanosine triphosphate (GTP)-binding proteins, which are closely related to intracellular calcium signaling. The underlying mechanism may involve the protein serine/threonine kinase pathway, cellular metabolism, and other cellular processes [131].

## 4. Outlook for Future Research

CMD poses a significant challenge to achieving clinical benefits in IHD, and effective interventions are currently limited, providing an opportunity for intervention with TCM. In recent years, several studies have demonstrated that TCM can improve CMD by protecting CMECs against various injuries. These interventions have shown structural improvements such as increased microvessel density and number, reduced microthrombosis, and inhibition of endothelial-to-mesenchymal transition. Functionally, TCM has been found to dilate coronary microvessels, alleviate microvascular permeability, and delay cellular senescence in CMECs. The mechanisms of action involve antioxidant, anti-apoptotic, anti-inflammatory effects, as well as regulation of energy metabolism, among others.

Considering that disease manifestation is often the result of multiple cell types within the same tissue or involving other tissues, co-culture systems have been used to mimic this condition, where different cell types share the same culture environment [132]. Recent studies have highlighted the ability of TCM to modulate the paracrine and autocrine secretion of CMECs, thereby influencing their own behavior and that of surrounding cells.

Enhancing the crosstalk between different cell types represents an important area for future research. Additionally, it has been observed that women have a higher risk of CMD compared to men [133], emphasizing the importance of studying gender differences in the pathophysiology of CMD [134]. Currently, preclinical studies rarely consider gender as a variable in their analyses [135]. Therefore, investigating the effects of CMD on CMECs in different genders may present a new research direction.

## 5. Limitations

The present study has several limitations that should be acknowledged. Firstly, the specific chemical structures of the active ingredients present in the Chinese medicines discussed in some articles remain unknown. Secondly, there are variations in the intervention methods used for compound Chinese medicines or the core components of Chinese medicines across different studies, leading to variations in the observed effects. Therefore, it is still necessary to establish standardized and uniform intervention methods to ensure consistency and comparability among studies.

## 6. Summary

This systematic review provides evidence that different injury models lead to distinct phenotypes in CMECs, and the intervention mechanisms of TCM also vary accordingly. Specifically, under ischemic or hypoxic injury conditions, TCM demonstrates its efficacy by promoting microangiogenesis, alleviating microvascular permeability, and inhibiting myocardial fibrosis. These effects are mediated through signaling pathways such as HIF-1α/VEGF, PI3K/AKT, and Snail. In the context of H/R injury, TCM exerts its benefits by inhibiting apoptosis, alleviating microvascular permeability, and dilating coronary microvessels, involving pathways such as ATM, PI3K/AKT, p70s6kT, p70s6kT/eNOS, and others. In the case of inflammatory injury, TCM acts by suppressing the release of inflammatory cytokines, regulating the expression of adhesion molecules, and reducing the formation of microthrombosis, through pathways such as MAPK, NF-κB, and JAK/STAT. Furthermore, under metabolic injury, angiotensin II, aging, and other pathological conditions, TCM demonstrates its efficacy by alleviating microvascular permeability, dilating coronary microvessels, and inhibiting inflammation and oxidative stress. The underlying signaling pathways involved include PPAR-α/Angptl4, H3K9ac/claudins, PI3K/AKT, among others. This systematic review provides a comprehensive overview of the effects of TCM on CMECs in various injury models and highlights the associated signaling pathway studies. These findings serve as a foundation for the application of TCM in the treatment of CMD.

## Abbreviations

4-ACGC: 4-O-(2-O-acetyl-6-O-p-coumaroyl-β-D-glucopyranosyl)-p-coumaric acid
A: Allicin
ADM: Adriamycin
AJs: Adherens junctions
AKT: Protein kinase B
ALDH: Aldehyde dehydrogenase
AMPK: Adenosine 5′-monophosphate activated protein kinase
AND: Andrographolide
Ang II: Angiotensin II
Angptl4: Angiopoietin-like 4
AP: Astragalus polysaccharide
ApoE: Apolipoprotein E
ARE: Antioxidant response element
ASIV: Astragaloside IV
ATP: Adenosine triphosphate
ATM: Ataxia telangiectasia mutated
bFGF: Basic fibroblast growth factor
BTE: Black tea extract
CBF: Coronary blood flow
CCM: Compound Chinese medicine
CFs: Cardiac fibroblasts
CFR: Coronary flow reserve
CHM: Chinese herbal medicines
CL: Carthamus tinctorius L. and Lepidium apetalum Willd
CMD: Coronary microvascular dysfunction
CMVD: Coronary microvascular disease
CMECs: Cardiac microvascular endothelial cells
COX-2: Cyclooxygenase-2
CTSS: Cathepsin S
CVB3: Coxsackievirus B3
CYP: Alpha-Cyperone
CYT: Caffeoylxanthiazonoside
DG: GeGen DanShen extract
DGBX: DangGuiBuXue Tang
Diosmetin-7-O-glucoside: Diosmetin-7-O-β-D-glucopyranoside
DLF: DanLou formula
Drp1: Dynamin-related protein 1
ECE-1: Endothelin-converting enzyme-1
EGb761: Ginkgo biloba extract
EndMT: Endothelial-to-mesenchymal transition
eNOS: Endothelial nitric oxide synthase
ER: Estrogen receptor
ERK: Extracellular signal-regulated kinase
ERS: Endoplasmic reticulum stress
FGF: Fibroblast growth factor
FoxO: Forkhead-Box Class O
GAS: Gastrodin
GTP: Guanosine triphosphate
H3K9ac: Histone H3 lysine 9 acetylation
H/R: Hypoxia/Reoxygenation
Hcy: Homocysteine
HF: Heart failure
HFD: High fat diet
HG: High glucose
HIF: Hypoxia-inducible factor
HO-1: Heme oxygenase-1
HSP27: Heat shock protein 27
I/R: Ischemia/reperfusion
IκB: Inhibitor of NF-κB
IGF1R: Insulin-like growth factor 1 receptor
IHD: Ischemic heart disease
IL: Interleukin
iNOS: Inducible nitric oxide synthase
IRS1: Insulin receptor substrate 1
JAK: Janus tyrosine kinase
JAM: Junctional adhesion molecule
KLF: Krüppel-like factor
L-NAME: N(ω)-nitro-L-arginine-methyl ester
LDL-C: Low-density lipoprotein cholesterol
Linc-ROR: Long intergenic non-coding RNA regulator of reprogramming
LPS: Lipopolysaccharide
MAPK: Mitogen-activated protein kinase
MBC: Major bioactive component
MDA: Malondialdehyde
MDG-1: A water-soluble beta-D-fructan from O. japonicus
MEK: Mitogen-activated protein kinase
MF: Myocardial fibrosis
Mff: mitochondrial fission factor
MHBFC: 17-Methoxyl-7-hydroxy-benzene-furanchalcone
MI: Myocardial infarction
MLC: Myosin light chain
MMP-9: Matrix metallopeptidase-9
mTOR: Mammalian target of rapamycin
NADPH: Nicotinamide adenine dinucleotide phosphate
NF-κB: Nuclear factor-kappa B
NLRP3: Nucleotide-binding oligomerization domain-like receptor protein 3
NO: Nitric oxide
NOS: Nitric oxide synthase
NR4A1: Nuclear receptor subfamily 4 group A member 1
Nrf2: Nuclear factor-erythroid 2 related factor 2
NRG-1: Neuregulin-1
NT: Nitrotyrosine
OA: Oroxylin A
OJ: Ophiopogon japonicus
P-IκBα: Phosphorylated inhibitor of kappa B alpha
P-p65: Phosphorylated p65
P70S6k1: 70 kDa ribosomal protein S6 kinase 1
PAI: Plasminogen activator inhibitor
PDGF: Platelet-derived growth factor
PECAM-1: Platelet-endothelial cell adhesion molecule-1
PER: Periplocin
PGE2: Prostaglandin E2
PGIS: Prostacyclin synthase
PHDs: Prolyl hydroxylase domain enzymes
PI3K: Phosphatidylinositol 3-kinase
PINK: Phosphatase and tensin homolog-induced putative kinase
PLCδ3: Phospholipase Cδ3
PMN: Polymorphonuclear leukocyte
PN: Panax Notoginseng
PPAR-α: Peroxisome proliferator-activated receptor α
PRISMA: Preferred Reporting Items for Systematic Reviews and Meta-Analyses
Pyk: Proline-rich tyrosine kinase
Q: Quercetin
QLQX: QiLiQiangXin
QSYQ: QiShenYiQi
Rb3: Ginsenoside-Rb3
RG: Radix Ginseng
Rg3: Ginsenoside-Rg3
RhoA: Ras homolog gene family member A
RIP3: Receptor interacting protein 3
RLC: Rhizoma Ligustici Chuanxiong
ROCK: Rho-associated protein kinase
ROS: Reactive oxygen species
RPS6KB1: Ribosomal protein S6 kinase B1
RS: Radix Scutellariae
RSV: Resveratrol
S1P: Sphingosine 1 phosphate
SA: Salidroside
SCU: Scutellarin
SL: ShenLian extract
SMF: ShenMai formula
SOD: Superoxide dismutase
SSNX: ShuangShenNingXin formula
STAT: Signal transducer and activator of transcription
STDP: SheXiangTongXin Dropping Pill
SXBX: SheXiangBaoXin Pill
Tan II A: Tanshinone II A
TCM: Traditional Chinese medicine
TF: Tissue factor
TGF: Transforming growth factor
TJs: Tight junctions
TMYX: TongMaiYangXin pill
TNF-α: Tumor necrosis factor-α
TXD: TianXiangDan
TXL: TongXinLuo
VA: Velvet Antler
VAP: Velvet antler proteins
VE: Vascular endothelial
VEGF: Vascular endothelial growth factor
VMC: Viral myocarditis
VSMCs: Vascular smooth muscle cells
XQT: XiangQiTang
ZO-1: Zona occludens-1.

## Acknowledgments

The authors would like to thank those who provided comments on the revision of this review.

## References

1. Chen W, Ni M, Huang H, Cong H, Fu X, Gao W, et al. Chinese expert consensus on the diagnosis and treatment of coronary microvascular diseases (2023 Edition). MedComm (2020). 2023;4(6):e438. pmid:38116064

2. Del Buono MG, Montone RA, Camilli M, Carbone S, Narula J, Lavie CJ, et al. Coronary Microvascular Dysfunction Across the Spectrum of Cardiovascular Diseases: JACC State-of-the-Art Review. J Am Coll Cardiol. 2021;78(13):1352–1371. pmid:34556322

3. Rush CJ, Berry C, Oldroyd KG, Rocchiccioli JP, Lindsay MM, Touyz RM, et al. Prevalence of Coronary Artery Disease and Coronary Microvascular Dysfunction in Patients With Heart Failure With Preserved Ejection Fraction. JAMA Cardiol. 2021;6(10):1130–1143. pmid:34160566

4. Bairey Merz CN, Pepine CJ, Shimokawa H, Berry C. Treatment of coronary microvascular dysfunction. Cardiovasc Res. 2020;116(4):856–870. pmid:32087007

5. Zhong L, Zhuang J, Jin Z, Chen Y, Chen B. Effect of Chinese medicine for promoting blood circulation on microvascular angina: A systematic review and meta-analysis. Am J Emerg Med. 2020;38(12):2681–2692. pmid:33046314

6. Yang Z, Lin S, Liu Y, Ren Q, Ge Z, Wang C, et al. Traditional chinese medicine in coronary microvascular disease. Front Pharmacol. 2022;13:929159. pmid:36003524

7. Deng J. Research progress on the molecular mechanism of coronary microvascular endothelial cell dysfunction. Int J Cardiol Heart Vasc. 2021;34:100777. pmid:33912653

8. Kaski JC, Crea F, Gersh BJ, Camici PG. Reappraisal of Ischemic Heart Disease. Circulation. 2018;138(14):1463–1480. pmid:30354347

9. Li JM, Mullen AM, Shah AM. Phenotypic properties and characteristics of superoxide production by mouse coronary microvascular endothelial cells. J Mol Cell Cardiol. 2001;33(6):1119–1131. pmid:11444917

10. Scarabelli T, Stephanou A, Rayment N, Pasini E, Comini L, Curello S, et al. Apoptosis of endothelial cells precedes myocyte cell apoptosis in ischemia/reperfusion injury. Circulation. 2001;104(3):253–256. pmid:11457740

11. McDouall RM, Batten P, McCormack A, Yacoub MH, Rose ML. MHC class II expression on human heart microvascular endothelial cells: exquisite sensitivity to interferon-gamma and natural killer cells. Transplantation. 1997;64(8):1175–1180. pmid:9355836

12. Alexander Y, Osto E, Schmidt-Trucksäss A, Shechter M, Trifunovic D, Duncker DJ, et al. Endothelial function in cardiovascular medicine: a consensus paper of the European Society of Cardiology Working Groups on Atherosclerosis and Vascular Biology, Aorta and Peripheral Vascular Diseases, Coronary Pathophysiology and Microcirculation, and Thrombosis. Cardiovasc Res. 2021;117(1):29–42. pmid:32282914

13. Knapp M, Tu X, Wu R. Vascular endothelial dysfunction, a major mediator in diabetic cardiomyopathy. Acta Pharmacol Sin. 2019;40(1):1–8. pmid:29867137

14. Crea F, Camici PG, Bairey Merz CN. Coronary microvascular dysfunction: an update. Eur Heart J. 2014;35(17):1101–1111. pmid:24366916

15. Page MJ, Moher D, Bossuyt PM, Boutron I, Hoffmann TC, Mulrow CD, et al. PRISMA 2020 explanation and elaboration: updated guidance and exemplars for reporting systematic reviews. Bmj. 2021;372:n160. pmid:33781993

16. Tousoulis D, Simopoulou C, Papageorgiou N, Oikonomou E, Hatzis G, Siasos G, et al. Endothelial dysfunction in conduit arteries and in microcirculation. Novel therapeutic approaches. Pharmacol Ther. 2014;144(3):253–267. pmid:24928320

17. Zhai S, Zhang XF, Lu F, Chen WG, He X, Zhang CF, et al. Chinese medicine GeGen-DanShen extract protects from myocardial ischemic injury through promoting angiogenesis via up-regulation of VEGF/VEGFR2 signaling pathway. J Ethnopharmacol. 2021;267:113475. WOS:000606371200001

18. Li Y, Wang Z, Mao M, Zhao M, Xiao X, Sun W, et al. Velvet Antler Mobilizes Endothelial Progenitor Cells to Promote Angiogenesis and Repair Vascular En dothelial Injury in Rats Following Myocardial Infarction. Front Physiol. 2018;9:1940. pmid:30705637

19. Hu GY, Yang P, Zeng Y, Zhang SY, Song J. Danggui Buxue decoction promotes angiogenesis by up-regulation of VEGFR(1/2) expressions and down-regulation of sVEGFR(1/2) expression in myocardial infarction rat. J Chin Med Assoc. 2018;81(1):37–46. WOS:000423780700008

20. Cui L, Liu Y, Hu Y, Dong J, Deng Q, Jiao B, et al. Shexiang Tongxin Dropping Pill alleviates M1 macrophage polarization-induced inflammation and endothelial dysfunction to reduce coronary microvascular dysfunction via the Dectin-1/Syk/IRF5 pathway. J Ethnopharmacol. 2023;316:116742.

21. Welt K, Fitzl G, Schaffranietz L. Myocardium-protective effects of Ginkgo biloba extract (EGb 761) in old rats against acute isobaric hypoxia. An electron microscopic morphometric study. II. Protection of microvascular endothelium. Exp Toxicol Pathol. 1996;48(1):81–86. pmid:8919274

22. Skuli N, Majmundar AJ, Krock BL, Mesquita RC, Mathew LK, Quinn ZL, et al. Endothelial HIF-2α regulates murine pathological angiogenesis and revascularization processes. J Clin Invest. 2012;122(4):1427–1443. pmid:22426208

23. Hirota K, Semenza GL. Regulation of angiogenesis by hypoxia-inducible factor 1. Crit Rev Oncol Hematol. 2006;59(1):15–26. pmid:16716598

24. Wang Y, Zhang J, Fu M, Wang J, Cui X, Song Y, et al. Qiliqiangxin Prescription Promotes Angiogenesis of Hypoxic Primary Rat Cardiac Microvascular Endothelial Cells via Regulating miR-21 Signaling. Curr Pharm Des. 2021;27(26):2966–2974. pmid:33019923

25. Pagé EL, Chan DA, Giaccia AJ, Levine M, Richard DE. Hypoxia-inducible factor-1alpha stabilization in nonhypoxic conditions: role of oxidation and intracellular ascorbate depletion. Mol Biol Cell. 2008;19(1):86–94. pmid:17942596

26. Wang Y, Han X, Fu M, Wang J, Song Y, Liu Y, et al. Qiliqiangxin attenuates hypoxia-induced injury in primary rat cardiac microvascular endothelial cells via promoting HIF-1α-dependent glycolysis. J Cell Mol Med. 2018;22(5):2791–2803. pmid:29502357

27. Xu Y, Li X, Liu X, Zhou M. Neuregulin-1/ErbB signaling and chronic heart failure. Adv Pharmacol. 2010;59:31–51. pmid:20933198

28. Wang JF, Zhou JM, Wang YY, Yang CJ, Fu MQ, Zhang JJ, et al. Qiliqiangxin protects against anoxic injury in cardiac microvascular endothelial cells via NRG-1/ErbB-PI3K/Akt/mTOR pathway. J Cell Mol Med. 2017;21(9):1905–1914. WOS:000408320500022

29. Li YN, Wu YL, Jia ZH, Qi JS. Interaction between COX-2 and iNOS aggravates vascular lesion and antagonistic effect of ginsenoside. J Ethnopharmacol. 2008;119(2):305–311. pmid:18694814

30. Tunctan B, Korkmaz B, Sari AN, Kacan M, Unsal D, Serin MS, et al. Contribution of iNOS/sGC/PKG pathway, COX-2, CYP4A1, and gp91(phox) to the protective effect of 5,14-HEDGE, a 20-HETE mimetic, against vasodilation, hypotension, tachycardia, and inflammation in a rat model of septic shock. Nitric Oxide. 2013;33:18–41. pmid:23684565

31. Li YN, Wang XJ, Li B, Liu K, Qi JS, Liu BH, et al. Tongxinluo inhibits cyclooxygenase-2, inducible nitric oxide synthase, hypoxia-inducible factor-2α/vascular endothelial growth factor to antagonize injury in hypoxia-stimulated cardiac microvascular endothelial cells. Chin Med J (Engl). 2015;128(8):1114–1120. pmid:25881609

32. Camacho M, Rodríguez C, Guadall A, Alcolea S, Orriols M, Escudero JR, et al. Hypoxia upregulates PGI-synthase and increases PGI₂ release in human vascular cells exposed to inflammatory stimuli. J Lipid Res. 2011;52(4):720–731. pmid:21296955

33. Liu B, Kuang L, Liu J. Bariatric surgery relieves type 2 diabetes and modulates inflammatory factors and coronary endothelium eNOS/iNOS expression in db/db mice. Can J Physiol Pharmacol. 2014;92(1):70–77. pmid:24383875

34. Wang X, Liu K, Li B, Li Y, Ye K, Qi J, et al. Macrophages Aggravate Hypoxia-Induced Cardiac Microvascular Endothelial Cell Injury via Peroxynitrite: Protection by Tongxinluo. Cell Commun Adhes. 2015;22(2-6):39–47. pmid:27001368

35. Robinson AS, Materna SC, Barnes RM, De Val S, Xu SM, Black BL. An arterial-specific enhancer of the human endothelin converting enzyme 1 (ECE1) gene is synergistically activated by Sox17, FoxC2, and Etv2. Dev Biol. 2014;395(2):379–389. pmid:25179465

36. Lee WL, Slutsky AS. Sepsis and endothelial permeability. New Engl J Med. 2010;363(7):689–691. pmid:20818861

37. Günzel D, Fromm M. Claudins and other tight junction proteins. Compr Physiol. 2012;2(3):1819–1852. pmid:23723025

38. Blaise S, Polena H, Vilgrain I. Soluble vascular endothelial-cadherin and auto-antibodies to human vascular endothelial-cadherin in human diseases: Two new biomarkers of endothelial dysfunction. Vasc Med. 2015;20(6):557–565. pmid:26129735

39. Nagafuchi A. Molecular architecture of adherens junctions. Curr Opin Cell Biol. 2001;13(5):600–603. pmid:11544029

40. Liu K, Wang XJ, Li YN, Li B, Qi JS, Zhang J, et al. Tongxinluo Reverses the Hypoxia-suppressed Claudin-9 in Cardiac Microvascular Endothelial Cells. Chin Med J (Engl). 2016;129(4):442–447. pmid:26879018

41. Zheng CY, Song LL, Wen JK, Li LM, Guo ZW, Zhou PP, et al. Tongxinluo (TXL), a Traditional Chinese Medicinal Compound, Improves Endothelial Function After Chronic Hypoxia Both In Vivo and In Vitro. J Cardiovasc Pharmacol. 2015;65(6):579–586. pmid:26065642

42. Lee SW, Won JY, Kim WJ, Lee J, Kim KH, Youn SW, et al. Snail as a potential target molecule in cardiac fibrosis: paracrine action of endothelial cells on fibroblasts through snail and CTGF axis. Mol Ther. 2013;21(9):1767–1777. pmid:23760445

43. Yin Y, Zhang Q, Zhao Q, Ding G, Wei C, Chang L, et al. Tongxinluo Attenuates Myocardiac Fibrosis after Acute Myocardial Infarction in Rats via Inhibition of Endothelial-to-Mesenchymal Transition. Biomed Res Int. 2019;2019:6595437. pmid:31317035

44. Zhou Z, Ma DF, Zhou Y, Zhang KK, Liu Y, Wang Z, et al. The Carthamus tinctorius L. and Lepidium apetalum Willd. Drug Pair Inhibits EndMT through the TGF beta 1/Snail Signaling Pathway in the Treatment of Myocardial Fibrosis. Evid Based Complement Alternat Med. 2023;2023:6018375. WOS:000919135800001

45. Dai GH, Liu N, Zhu JW, Yao J, Yang C, Ma PZ, et al. Qi-Shen-Yi-Qi Dripping Pills Promote Angiogenesis of Ischemic Cardiac Microvascular Endothelial Cells by Regulating MicroRNA-223-3p Expression. Evid Based Complement Alternat Med. 2016;2016:5057328. WOS:000370705200001

46. Wang SO, Lin X, Wang LY, Ruan KF, Feng Y, Li XY. A polysaccharides MDG-1 augments survival in the ischemic heart by inducing S1P release and S1P(1) expression. Int J Biol Macromol. 2012;50(3):734–740. WOS:000302664900040

47. Bastaki M, Nelli EE, Dell’Era P, Rusnati M, Molinari-Tosatti MP, Parolini S, et al. Basic fibroblast growth factor-induced angiogenic phenotype in mouse endothelium. A study of aortic and microvascular endothelial cell lines. Arterioscler Thromb Vasc Biol. 1997;17(3):454–464. pmid:9102163

48. Wang SO, Zhang Z, Lin X, Xu DS, Feng Y, Ding K. A polysaccharide, MDG-1, induces S1P(1) and bFGF expression and augments survival and angiogenesis in the ischemic heart. Glycobiology. 2010;20(4):473–484. WOS:000275567900008

49. Hu J, Zhao Y, Wu Y, Yang K, Hu K, Sun A, et al. Shexiang Baoxin Pill Attenuates Ischemic Injury by Promoting Angiogenesis by Activation of Aldehyde Dehydrogenase 2. J Cardiovasc Pharmacol. 2021;77(3):408–417. pmid:33662981

50. Xiao X, Xu SQ, Li L, Mao M, Wang JP, Li YJ, et al. The Effect of Velvet Antler Proteins on Cardiac Microvascular Endothelial Cells Challenged with Ischemia-Hypoxia. Front Pharmacol. 2017;8:601. WOS:000409379900001

51. Zhao N, Liu YY, Wang F, Hu BH, Sun K, Chang X, et al. Cardiotonic pills, a compound Chinese medicine, protects ischemia-reperfusion-induced microcirculatory disturbance and myocardial damage in rats. Am J Physiol Heart Circ Physiol. 2010;298(4):H1166–1176. pmid:20118406

52. Chen R, Chen T, Wang TQ, Dai XD, Meng K, Zhang SY, et al. Tongmai Yangxin pill reduces myocardial no-reflow by regulating apoptosis and activating PI3K/Akt/eNOS pathway. J Ethnopharmacol. 2020;261:113069. WOS:000566339300005

53. Dai X, Chen R, Chen T, Yan H, Wang Y, Zhou K, et al. Danlou Fang (丹蒌方) reduces microvascular obstruction through the endothelial/inducible nitric oxide synthase pathway in a rat model. J Tradit Chin Med. 2021;41(2):246–253.

54. Cui H, Yang Y, Li X, Zong W, Li Q. Resveratrol regulates paracrine function of cardiac microvascular endothelial cells under hypoxia/reoxygenation condition. Pharmazie. 2022;77(6):179–185. pmid:35751162

55. Shi MN, Liu YT, Feng LX, Cui YB, Chen YJ, Wang P, et al. Protective Effects of Scutellarin on Human Cardiac Microvascular Endothelial Cells against Hypoxia-Reoxygenation Injury and Its Possible Target-Related Proteins. Evid Based Complement Alternat Med. 2015;2015:278014. WOS:000364049100001

56. Mehta D, Malik AB. Signaling mechanisms regulating endothelial permeability. Physiol Rev. 2006;86(1):279–367. pmid:16371600

57. Pan CS, Yan L, Lin SQ, He K, Cui YC, Liu YY, et al. QiShenYiQi Pills Attenuates Ischemia/Reperfusion-Induced Cardiac Microvascular Hyperpermeability Implicating Src/Caveolin-1 and RhoA/ROCK/MLC Signaling. Front Physiol. 2021;12:753761. pmid:34975519

58. He K, Yan L, Lin SQ, Liu YY, Hu BH, Chang X, et al. Implication of IGF1R signaling in the protective effect of Astragaloside IV on ischemia and reperfusion-induced cardiac microvascular endothelial hyperpermeability. Phytomedicine. 2022;100:154045. pmid:35338991

59. Lefer AM, Lefer DJ. The role of nitric oxide and cell adhesion molecules on the microcirculation in ischaemia-reperfusion. Cardiovasc Res. 1996;32(4):743–751. pmid:8915192

60. Hai-Yan Z, Yong-Hong G, Zhi-Yao W, Bing X, Ai-Ming W, Yan-Wei X, et al. Astragalus Polysaccharide Suppresses the Expression of Adhesion Molecules through the Regulation of the p38 MAPK Signaling Pathway in Human Cardiac Microvascular Endothelial Cells after Ischemia-Reperfusion Injury. Evid Based Complement Alternat Med. 2013;2013:280493. pmid:24302961

61. Li Q, Cui HH, Yang YJ, Li XD, Chen GH, Tian XQ, et al. Quantitative Proteomics Analysis of Ischemia/Reperfusion Injury-Modulated Proteins in Cardiac Microvascular Endothelial Cells and the Protective Role of Tongxinluo. Cell Physiol Biochem. 2017;41(4):1503–1518. WOS:000401163100021

62. Cui H, Li N, Li X, Qi K, Li Q, Jin C, et al. Tongxinluo modulates cytokine secretion by cardiac microvascular endothelial cells in ischemia/reperfusion injury. Am J Transl Res. 2016;8(10):4370–4381.

63. Chen G, Xu C, Gillette TG, Huang T, Huang P, Li Q, et al. Cardiomyocyte-derived small extracellular vesicles can signal eNOS activation in cardiac microvascular endothelial cells to protect against Ischemia/Reperfusion injury. Theranostics. 2020;10(25):11754–11774. pmid:33052245

64. Ginsberg MD. Expanding the concept of neuroprotection for acute ischemic stroke: The pivotal roles of reperfusion and the collateral circulation. Prog Neurobiol. 2016;145–146:46-77. pmid:27637159

65. Cui HH, Li XD, Li N, Qi K, Li Q, Jin C, et al. Induction of Autophagy by Tongxinluo Through the MEK/ERK Pathway Protects Human Cardiac Microvascular Endothelial Cells From Hypoxia/Reoxygenation Injury. J Cardiovasc Pharmacol. 2014;64(2):180–190. WOS:000340727000009

66. Li JJ, Wang YJ, Wang CM, Li YJ, Yang Q, Cai WY, et al. Shenlian extract decreases mitochondrial autophagy to regulate mitochondrial function in microvascular to alleviate coronary artery no-reflow. Phytother Res. 2023;37(5):1864–1882. pmid:36740450

67. Liu Z, Han X, You Y, Xin G, Li L, Gao J, et al. Shuangshen ningxin formula attenuates cardiac microvascular ischemia/reperfusion injury through improving mitochondrial function. J Ethnopharmacol. 2024;323:117690. pmid:38195019

68. Sheng J, Li H, Dai Q, Lu C, Xu M, Zhang J, et al. NR4A1 Promotes Diabetic Nephropathy by Activating Mff-Mediated Mitochondrial Fission and Suppressing Parkin-Mediated Mitophagy. Cell Physiol Biochem. 2018;48(4):1675–1693. pmid:30077998

69. Wang J, Toan S, Zhou H. Mitochondrial quality control in cardiac microvascular ischemia-reperfusion injury: New insights into the mechanisms and therapeutic potentials. Pharmacol Res. 2020;156:104771. pmid:32234339

70. Zhang Z, Li X, He J, Wang S, Wang J, Liu J, et al. Molecular mechanisms of endothelial dysfunction in coronary microcirculation dysfunction. J Thromb Thrombolysis. 2023;56(3):388–397. pmid:37466848

71. Zhang C, Wang DF, Zhang Z, Han D, Yang K. EGb 761 Protects Cardiac Microvascular Endothelial Cells against Hypoxia/Reoxygenation Injury and Exerts Inhibitory Effect on the ATM Pathway. J Microbiol Biotechnol. 2017;27(3):584–590. pmid:27974731

72. Wang F, Miao M, Xia H, Yang LG, Wang SK, Sun GJ. Antioxidant activities of aqueous extracts from 12 Chinese edible flowers in vitro and in vivo. Food Nutr Res. 2017;61(1):1265324. pmid:28326000

73. Sun W, Lu H, Lyu L, Yang P, Lin Z, Li L, et al. Gastrodin ameliorates microvascular reperfusion injury-induced pyroptosis by regulating the NLRP3/caspase-1 pathway. J Physiol Biochem. 2019;75(4):531–547. pmid:31440987

74. Zanatta E, Colombo C, D’Amico G, d’Humières T, Dal Lin C, Tona F. Inflammation and Coronary Microvascular Dysfunction in Autoimmune Rheumatic Diseases. Int J Mol Sci. 2019;20(22):5563. pmid:31703406

75. Pober JS, Sessa WC. Evolving functions of endothelial cells in inflammation. Nat Rev Immunol. 2007;7(10):803–815. pmid:17893694

76. Zhu JQ, Liang YB, Yue SQ, Fan GW, Zhang H, Zhang M. Combination of Panaxadiol and Panaxatriol Type Saponins and Ophioponins From Shenmai Formula Attenuates Lipopolysaccharide-induced Inflammatory Injury in Cardiac Microvascular Endothelial Cells by Blocking NF-kappa B Pathway. J Cardiovasc Pharmacol. 2017;69(3):140–146. WOS:000395798500003

77. Han S, Wu H, Li W, Gao P. Protective effects of genistein in homocysteine-induced endothelial cell inflammatory injury. Mol Cell Biochem. 2015;403(1-2):43–49. pmid:25626894

78. Zhang H, Park Y, Wu J, Chen X, Lee S, Yang J, et al. Role of TNF-alpha in vascular dysfunction. Clin Sci (Lond). 2009;116(3):219–230. pmid:19118493

79. Collins T, Read MA, Neish AS, Whitley MZ, Thanos D, Maniatis T. Transcriptional regulation of endothelial cell adhesion molecules: NF-kappa B and cytokine-inducible enhancers. FASEB J. 1995;9(10):899–909. pmid:7542214

80. Pan LL, Dai M. Paeonol from Paeonia suffruticosa prevents TNF-alpha-induced monocytic cell adhesion to rat aortic endothelial cells by suppression of VCAM-1 expression. Phytomedicine. 2009;16(11):1027–1032. pmid:19541467

81. Yang Y, Yu K, Zhang YM. The Cardioprotective Effects of 4-O-(2″-O-acetyl-6″-O-P-coumaroyl-β-D-glucopyranosyl)-P-coumaric Acid (4-ACGC) on Chronic Heart Failure. Iran J Pharm Res. 2018;17(2):593–600. pmid:29881417

82. Li R, Dong Z, Zhuang X, Liu R, Yan F, Chen Y, et al. Salidroside prevents tumor necrosis factor-α-induced vascular inflammation by blocking mitogen-activated protein kinase and NF-κB signaling activation. Exp Ther Med. 2019;18(5):4137–4143. pmid:31656544

83. Sawuer G, Ma XK, Zhang YJ, Zhang XM, Ainiwaer Z, An DQ. Tianxiangdan Improves Coronary Microvascular Dysfunction in Rats by Inhibiting Microvascular Inflammation via Nrf2 Activation. Evid Based Complement Alternat Med. 2021;2021:4414784. WOS:000788538200003

84. Bagheri F, Khori V, Alizadeh AM, Khalighfard S, Khodayari S, Khodayari H. Reactive oxygen species-mediated cardiac-reperfusion injury: Mechanisms and therapies. Life Sci. 2016;165:43–55. pmid:27667751

85. Touyz RM, Yao G, Schiffrin EL. c-Src induces phosphorylation and translocation of p47phox: role in superoxide generation by angiotensin II in human vascular smooth muscle cells. Arterioscler Thromb Vasc Biol. 2003;23(6):981–987. pmid:12663375

86. Wu XL, Zheng B, Jin LS, Zhang RN, Ming H, Yang Z, et al. Chinese medicine Tongxinluo reduces atherosclerotic lesion by attenuating oxidative stress and inflammation in microvascular endothelial cells. Int J Clin Exp Pathol. 2015;8(6):6323–6333. WOS:000359277700031

87. Yang B, Wang F, Cao H, Liu G, Zhang Y, Yan P, et al. Caffeoylxanthiazonoside exerts cardioprotective effects during chronic heart failure via inhibition of inflammatory responses in cardiac cells. Exp Ther Med. 2017;14(5):4224–4230. pmid:29104638

88. Dellinger RP. Inflammation and coagulation: implications for the septic patient. Clin Infect Dis. 2003;36(10):1259–1265. pmid:12746771

89. Aiuto LT, Barone SR, Cohen PS, Boxer RA. Recombinant tissue plasminogen activator restores perfusion in meningococcal purpura fulminans. Crit Care Med. 1997;25(6):1079–1082. pmid:9201064

90. Shi RJ, Simpson-Haidaris PJ, Lerner NB, Marder VJ, Silverman DJ, Sporn LA. Transcriptional regulation of endothelial cell tissue factor expression during Rickettsia rickettsii infection: involvement of the transcription factor NF-kappaB. Infect Immun. 1998;66(3):1070–1075. pmid:9488397

91. He CL, Yi PF, Fan QJ, Shen HQ, Jiang XL, Qin QQ, et al. Xiang-Qi-Tang and its active components exhibit anti-inflammatory and anticoagulant properties by inhibiting MAPK and NF-kappa B signaling pathways in LPS-treated rat cardiac microvascular endothelial cells. Immunopharmacol Immunotoxicol. 2013;35(2):215–224. WOS:000316125800002

92. Frank PG, Lee H, Park DS, Tandon NN, Scherer PE, Lisanti MP. Genetic ablation of caveolin-1 confers protection against atherosclerosis. Arterioscler Thromb Vasc Biol. 2004;24(1):98–105. pmid:14563650

93. Feng B, Zhang Q, Wang X, Sun X, Mu X, Dong H. Effect of Andrographolide on Gene Expression Profile and Intracellular Calcium in Primary Rat Myocardium Microvascular Endothelial Cells. J Cardiovasc Pharmacol. 2017;70(6):369–381. pmid:29112590

94. Lakota K, Mrak-Poljsak K, Rozman B, Sodin-Semrl S. Increased responsiveness of human coronary artery endothelial cells in inflammation and coagulation. Mediators Inflamm. 2009;2009:146872. pmid:20107610

95. Zhang Y, Zhu MD, Zhang FG, Zhang SQ, Du WX, Xiao XF. Integrating Pharmacokinetics Study, Network Analysis, and Experimental Validation to Uncover the Mechanism of Qiliqiangxin Capsule Against Chronic Heart Failure. Front Pharmacol. 2019;10:1046. WOS:000486436900001

96. Vergès B. Clinical interest of PPARs ligands. Diabetes Metab. 2004;30(1):7–12. pmid:15029092

97. Lakka HM, Laaksonen DE, Lakka TA, Niskanen LK, Kumpusalo E, Tuomilehto J, et al. The metabolic syndrome and total and cardiovascular disease mortality in middle-aged men. Jama. 2002;288(21):2709–2716. pmid:12460094

98. Khazaei M, Tahergorabi Z. Ghrelin did not change coronary angiogenesis in diet-induced obese mice. Cell Mol Biol (Noisy-le-grand). 2017;63(2):96–99. pmid:28364789

99. Yu S, Kim SR, Jiang K, Ogrodnik M, Zhu XY, Ferguson CM, et al. Quercetin Reverses Cardiac Systolic Dysfunction in Mice Fed with a High-Fat Diet: Role of Angiogenesis. Oxid Med Cell Longev. 2021;2021:8875729. pmid:33688395

100. Rosenfeld ME, Polinsky P, Virmani R, Kauser K, Rubanyi G, Schwartz SM. Advanced atherosclerotic lesions in the innominate artery of the ApoE knockout mouse. Arterioscler Thromb Vasc Biol. 2000;20(12):2587–2592. pmid:11116057

101. Pechánová O, Bernátová I, Pelouch V, Babál P. L-NAME-induced protein remodeling and fibrosis in the rat heart. Physiol Res. 1999;48(5):353–362. pmid:10625224

102. Huang Y, Zhang K, Liu M, Su J, Qin X, Wang X, et al. An herbal preparation ameliorates heart failure with preserved ejection fraction by alleviating microvascular endothelial inflammation and activating NO-cGMP-PKG pathway. Phytomedicine. 2021;91:153633. pmid:34320423

103. Li B, Li YN, Liu K, Wang XJ, Qi JS, Wang BY, et al. High glucose decreases claudins-5 and-11 in cardiac microvascular endothelial cells: Antagonistic effects of tongxinluo. Endocr Res. 2017;42(1):15–21. WOS:000394615200003

104. Galaup A, Gomez E, Souktani R, Durand M, Cazes A, Monnot C, et al. Protection against myocardial infarction and no-reflow through preservation of vascular integrity by angiopoietin-like 4. Circulation. 2012;125(1):140–149. pmid:22086875

105. Qi K, Yang YJ, Geng YJ, Cui HH, Li XD, Jin C, et al. Tongxinluo attenuates oxygen-glucose-serum deprivation/restoration-induced endothelial barrier breakdown via peroxisome proliferator activated receptor-alpha/angiopoietin-like 4 pathway in high glucose-incubated human cardiac microvascular endothelial cells. Medicine. 2020;99(34):e21821. WOS:000579455900078

106. Qi K, Li X, Geng Y, Cui H, Jin C, Wang P, et al. Tongxinluo attenuates reperfusion injury in diabetic hearts by angiopoietin-like 4-mediated protection of endothelial barrier integrity via PPAR-α pathway. PLoS One. 2018;13(6):e0198403. pmid:29912977

107. Lemos Caldas FR, Rocha Leite IM, Tavarez Filgueiras AB, de Figueiredo Júnior IL, Gomes Marques de Sousa TA, Martins PR, et al. Mitochondrial ATP-sensitive potassium channel opening inhibits isoproterenol-induced cardiac hypertrophy by preventing oxidative damage. J Cardiovasc Pharmacol. 2015;65(4):393–397. pmid:25850726

108. Yang K, Wang C, Nie L, Zhao X, Gu J, Guan X, et al. Klotho Protects Against Indoxyl Sulphate-Induced Myocardial Hypertrophy. J Am Soc Nephrol. 2015;26(10):2434–2446. pmid:25804281

109. Zeng H, Chen JX. Microvascular Rarefaction and Heart Failure With Preserved Ejection Fraction. Front Cardiovasc Med. 2019;6:15. pmid:30873415

110. Shi P, Cao Y, Gao J, Fu B, Ren J, Ba L, et al. Allicin improves the function of cardiac microvascular endothelial cells by increasing PECAM-1 in rats with cardiac hypertrophy. Phytomedicine. 2018;51:241–254. pmid:30466623

111. Guo X, Zhang Y, Lu C, Qu F, Jiang X. Protective effect of hyperoside on heart failure rats via attenuating myocardial apoptosis and inducing autophagy. Biosci Biotechnol Biochem. 2020;84(4):714–724. pmid:31797747

112. Li F, Wang J, Song Y, Shen D, Zhao Y, Li C, et al. Qiliqiangxin alleviates Ang II-induced CMECs apoptosis by downregulating autophagy via the ErbB2-AKT-FoxO3a axis. Life Sci. 2021;273:119239. pmid:33652033

113. Sengupta A, Molkentin JD, Yutzey KE. FoxO transcription factors promote autophagy in cardiomyocytes. J Biol Chem. 2009;284(41):28319–28331. pmid:19696026

114. Li LM, Zheng B, Zhang RN, Jin LS, Zheng CY, Wang C, et al. Chinese medicine Tongxinluo increases tight junction protein levels by inducing KLF5 expression in microvascular endothelial cells. Cell Biochem Funct. 2015;33(4):226–234. WOS:000356493500009

115. Rodríguez-Mañas L, El-Assar M, Vallejo S, López-Dóriga P, Solís J, Petidier R, et al. Endothelial dysfunction in aged humans is related with oxidative stress and vascular inflammation. Aging Cell. 2009;8(3):226–238. pmid:19245678

116. Wang Q, Yang J, Lei Y, Xiu C, Huo Y, Shi H. “Effects of extracts from Renshen (Radix Ginseng), Sanqi (Radix Notoginseng), and Chuanxiong (Rhizoma Chuanxiong) on F-actin in senescent microvascular endothelial cells”. J Tradit Chin Med. 2020;40(3):376–385. pmid:32506850

117. Pollack A, Kontorovich AR, Fuster V, Dec GW. Viral myocarditis--diagnosis, treatment options, and current controversies. Nat Rev Cardiol. 2015;12(11):670–680. pmid:26194549

118. Cai Z, Yang M, Huang L, Cheng L, Li H, Chen C. [Dynamic changes between osteopontin and collagen I expression in viral myocarditis mice]. Zhong Nan Da Xue Xue Bao Yi Xue Ban. 2012;37(3):271–277. pmid:22561508

119. Frangogiannis NG. Cardiac fibrosis. Cardiovasc Res. 2021;117(6):1450–1488. pmid:33135058

120. Zeisberg EM, Tarnavski O, Zeisberg M, Dorfman AL, McMullen JR, Gustafsson E, et al. Endothelial-to-mesenchymal transition contributes to cardiac fibrosis. Nat Med. 2007;13(8):952–961. pmid:17660828

121. Yang L, Liu Q, Yu Y, Xu H, Chen S, Shi S. Ginsenoside-Rb3 inhibits endothelial-mesenchymal transition of cardiac microvascular endothelial cells. Herz. 2019;44(1):60–68. pmid:28983639

122. Wang X, Chen L, Wang T, Jiang X, Zhang H, Li P, et al. Ginsenoside Rg3 antagonizes adriamycin-induced cardiotoxicity by improving endothelial dysfunction from oxidative stress via upregulating the Nrf2-ARE pathway through the activation of akt. Phytomedicine. 2015;22(10):875–884. pmid:26321736

123. Jian J, Huang JC, Jiao Y, Tan HD, Huang RB. Isolation and preparation of chalcone compounds from tuber of Millettia pulchra var. laxior by pre-HPLC. Chinese Traditional and Herbal Drugs. 2011;42(7):1313–1316.

124. Ye FX, He JH, Wu XM, Xie JX, Chen HL, Tang XJ, et al. The regulatory mechanisms of Yulangsan MHBFC reversing cardiac remodeling in rats based on eNOS-NO signaling pathway. Biomed Pharmacother. 2019;117:109141. WOS:000477804500117

125. Li A, Sun A, Liu R, Zhang Y, Cui J. An efficient preparative procedure for main flavonoids from the peel of Trichosanthes kirilowii Maxim. using polyamide resin followed by semi-preparative high performance liquid chromatography. J Chromatogr B Analyt Technol Biomed Life Sci. 2014;965:150–157. pmid:25023212

126. Wang H, Zhang X, Liu Y, Zhang Y, Wang Y, Peng Y, et al. Diosmetin-7-O-β-D-glucopyranoside suppresses endothelial-mesenchymal transformation through endoplasmic reticulum stress in cardiac fibrosis. Clin Exp Pharmacol Physiol. 2023;50(10):789–805. pmid:37430476

127. Lan S, Yi F, Shuang L, Chenjie W, Zheng XW. Chemical constituents from the fibrous root of Ophiopogon japonicus, and their effect on tube formation in human myocardial microvascular endothelial cells. Fitoterapia. 2013;85:57–63. pmid:23274777

128. Hu F, Chan JYW, Koon CM, Fung KP. Angiogenic Effects of Danshen and Gegen Decoction on Human Endothelial Cells and Zebrafish Embryos. Am J Chin Med. 2013;41(4):887–900. WOS:000322444500011

129. Qu J, Liu F, Zhang X, Wang J. Oroxylin A Reduces Vasoconstriction in Rat Aortic Rings through Promoting NO Production and NOS Protein Expression via Estrogen Receptor Signal Pathway. Evid Based Complement Alternat Med. 2020;2020:9257950. pmid:32082399

130. Fan G, Zhu Y, Guo H, Wang X, Wang H, Gao X. Direct vasorelaxation by a novel phytoestrogen tanshinone IIA is mediated by nongenomic action of estrogen receptor through endothelial nitric oxide synthase activation and calcium mobilization. J Cardiovasc Pharmacol. 2011;57(3):340–347. pmid:21383591

131. Wang XY, Gao XM, Liu H, Zhang H, Liu Y, Jiang M, et al. Gene expression profiling of the proliferative effect of periplocin on mouse cardiac microvascular endothelial cells. Chin J Integr Med. 2010;16(1):33–40. pmid:20131034

132. Lippi M, Stadiotti I, Pompilio G, Sommariva E. Human Cell Modeling for Cardiovascular Diseases. Int J Mol Sci. 2020;21(17):6388. pmid:32887493

133. Mileva N, Nagumo S, Mizukami T, Sonck J, Berry C, Gallinoro E, et al. Prevalence of Coronary Microvascular Disease and Coronary Vasospasm in Patients With Nonobstructive Coronary Artery Disease: Systematic Review and Meta-Analysis. J Am Heart Assoc. 2022;11(7):e023207. pmid:35301851

134. Maselli A, Matarrese P, Straface E, Canu S, Franconi F, Malorni W. Cell sex: a new look at cell fate studies. FASEB J. 2009;23(4):978–984. pmid:19074513

135. Allegra S, Chiara F, Di Grazia D, Gaspari M, De Francia S. Evaluation of Sex Differences in Preclinical Pharmacology Research: How Far Is Left to Go? Pharmaceuticals (Basel). 2023;16(6):786. pmid:37375734

